# Single-Component Photo-Crosslinked Lipoic Acid Functionalized Mucin Hydrogels: Dynamic Mucus Mimetics for Antiviral Protection

**DOI:** 10.64898/2026.11.08.765147

**Authors:** Ali Abbas, Hung-Yi Lin, Ayesha Iftikhar Cheema, Jui-Hong Weng, Jung-Ren Huang, Hung-Wei Hsu, I-Hsuan Wang, Chia-Fu Chou

## Abstract

Native mucus protects epithelial surfaces through viscoelastic networks of disulfide-linked mucins. However, commercially available mucins often fail to reconstitute comparable mechanical and barrier properties, while synthetic and hybrid mucus mimetics require polymeric crosslinkers that may *alter* mucus-like composition. Inspired by the native architecture, we directly functionalized porcine gastric and bovine submaxillary mucins with *α*-lipoic acid to develop a single-component, photo-crosslinkable hydrogels. Upon UV irradiation (365 nm), the modified mucins rapidly form disulfide-crosslinked networks without added photo-initiator/ polymeric crosslinkers, two orders of magnitude faster as compared with PEG-SH approaches. The resulting mucin-lipoic acid (MUC-LA) hydrogels exhibit mucus-relevant properties, including tunable viscoelasticity, redox responsiveness, self-healing, and size-dependent restrictions on nanoparticle transport. In transwell-based infection models, MUC-LA hydrogels (2-3%) mimicking the healthy airway mucus (G′ ≈ 10-20 Pa) suppress epithelial infection by SARS-CoV-2 and influenza A virus.

Crosslinked MUC-LA provides greater protection against SARS-CoV-2 than the non-crosslinked, whereas both formulations suppress influenza A infection, suggesting distinct contributions from network formation and mucin-associated interactions. These findings establish a dynamic, mucusmimetic platform integrating rapid photo-gelation, controllable mechanics, selective transport, and antiviral barrier functionality. MUC-LA hydrogels offer opportunities to investigate mucosal transport and host-pathogen interactions, with potential applications in drug delivery, and as biomaterials.

## 1 Introduction

Mucus is hydrated, viscoelastic barrier that covers epithelial surfaces and protects against environmental particles, toxins and microbial pathogens, including bacteria and viruses [1–2]. These protective properties arise mainly from mucins, heavily O-glycosylated proteins such as MUC2, MUC5AC and MUC5B, which forms heterogenous disulfide dominated network with characteristic dimensions spanning the nanometer scale [3–5]. Cysteine-rich domains facilitate intermolecular disulfide bonding and mucin polymerization while physical entanglements and noncovalent interactions contribute to the dynamic viscoelastic properties of mucus [6–7]. The balance between network mechanics and selective transport is essential for effective barrier function [8], as illustrated by impaired mucociliary clearance in diseases such as cystic fibrosis [9–10]. Reproducing this balance remains difficult in current *in vitro* models, creating a need for controllable mucus mimetics that reproduce both the rheological and barrier properties of native mucus [11].

Native mucus itself is difficult to study because of sample-to-sample variability and the complexity of isolation and purification procedures [12]. Commercial mucins such as porcine gastric mucin (PGM) and bovine submaxillary mucin (BSM) are widely used but often show limited gel-forming ability under physiological conditions due to loss of their native structure while processing [13–16]. Various approaches have been used to isolate mucins from biological tissues or cell-derived sources while preserving their gel-forming properties. However, these methods remain complex and low-yielding, which limits their scalability and broader applications [17–19].

To overcome these limitations, several synthetic and hybrid mucus mimetics have been explored. Bej et al., developed mucus-inspired hydrogels based on synthetic oligoglycerol (OG) polymers and multi-arm thiolated polyethylene glycol (PEG-SH) crosslinkers to form a disulfide network with self-healing and antiviral barrier properties [20]. Other systems have used synthetic mucin-like polymers or lignin-based hydrogels to reproduce selected aspects of mucus transport and barrier function [21–22]. Commercial mucins have also been incorporated into hybrid systems through external crosslinkers or covalent modification, including genipin-mediated and PEG-thiolbased disulfide networks [11, 23–26]. These approaches demonstrate the potential of mucusmimetic materials. However, most systems still rely on external crosslinking strategies or synthetic components, making it difficult to retain mucin as the main structural and functional component of the network. To the best of our knowledge, commercially available mucins have not yet been engineered to directly form a dynamic disulfide-crosslinked network without the addition of any external polymer/cross linkers.

Here, we address this gap by developing a mucin-dominated hydrogel in which commercially available PGM and BSM are directly functionalized with α-lipoic acid (LA), a cyclic disulfidecontaining molecule that introduces photo-responsive and redox-active sites while retaining mucin as the main network-forming component [27–28]. The strained five-membered 1,2-dithiolane ring of LA undergoes photo-induced ring opening under UV irradiation, enabling intermolecular disulfide network formation without the need for an added photo-initiator [29–30]. This strategy provides temporal control over gelation while avoiding the use of additional polymers. **Figure 1** summarizes the overall design of the mucin-lipoic acid (MUC-LA) platform, from LA functionalization and UV-triggered gelation to self-healing and barrier function.

**Figure 1.**
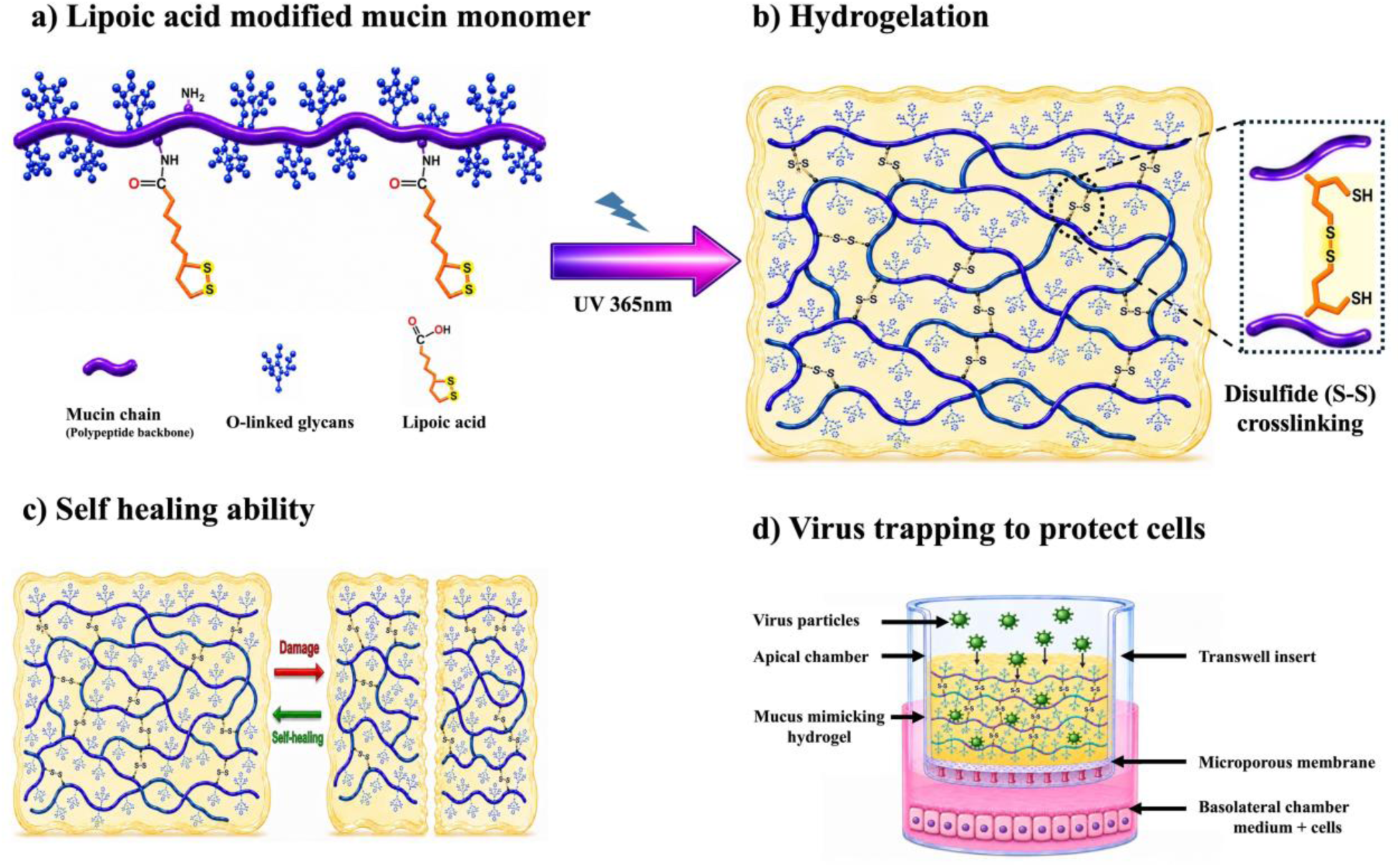
Schematic of the lipoic acid-modified mucin hydrogel and its characterization. (a) Lipoic acid-functionalized mucin monomer. (b) UV irradiation at 365 nm induces disulfide crosslinking and hydrogel formation. (c) Dynamic disulfide bonds enable self-healing after damage. (d) The hydrogel restricts viral particles transport and protects underlying cells from infection.

In this study, we developed a mucin-based MUC-LA hydrogel and investigated its photo-triggered gelation and dynamic disulfide network formation. Bulk rheology was used to characterize its mechanical properties, while redox-responsive and cyclic strain measurements were used to evaluate network adaptability and self-healing. Single-particle tracking was used to study sizedependent nanoparticle transport restriction within the hydrogel. Finally, transwell-based infection assays with SARS-CoV-2 and influenza A virus were used to evaluate the ability of the hydrogel to limit viral access to underlying epithelial cells. Together, these studies establish MUC-LA as a reproducible mucus-mimetic platform for investigating particle transport restriction, host-pathogen interactions and evaluating mucosal drug delivery.

## 2 Results and Discussion

### 2.1 Synthesis and Characterization of Lipoic Acid Conjugated Mucin

To generate a mucin (MUC) based hydrogel with dynamic disulfide crosslinking, lipoic acid (LA) was conjugated to accessible amine (-NH_2_) groups on the mucin backbone through EDC/NHS coupling reactions. This modification introduced dithiolane-containing side groups which can participate in photo-triggered disulfide network formation upon UV irradiation, *without* the addition of any photo-initiator or external crosslinker. The overall strategy is illustrated in **Figure 2a**.

**Figure 2.**
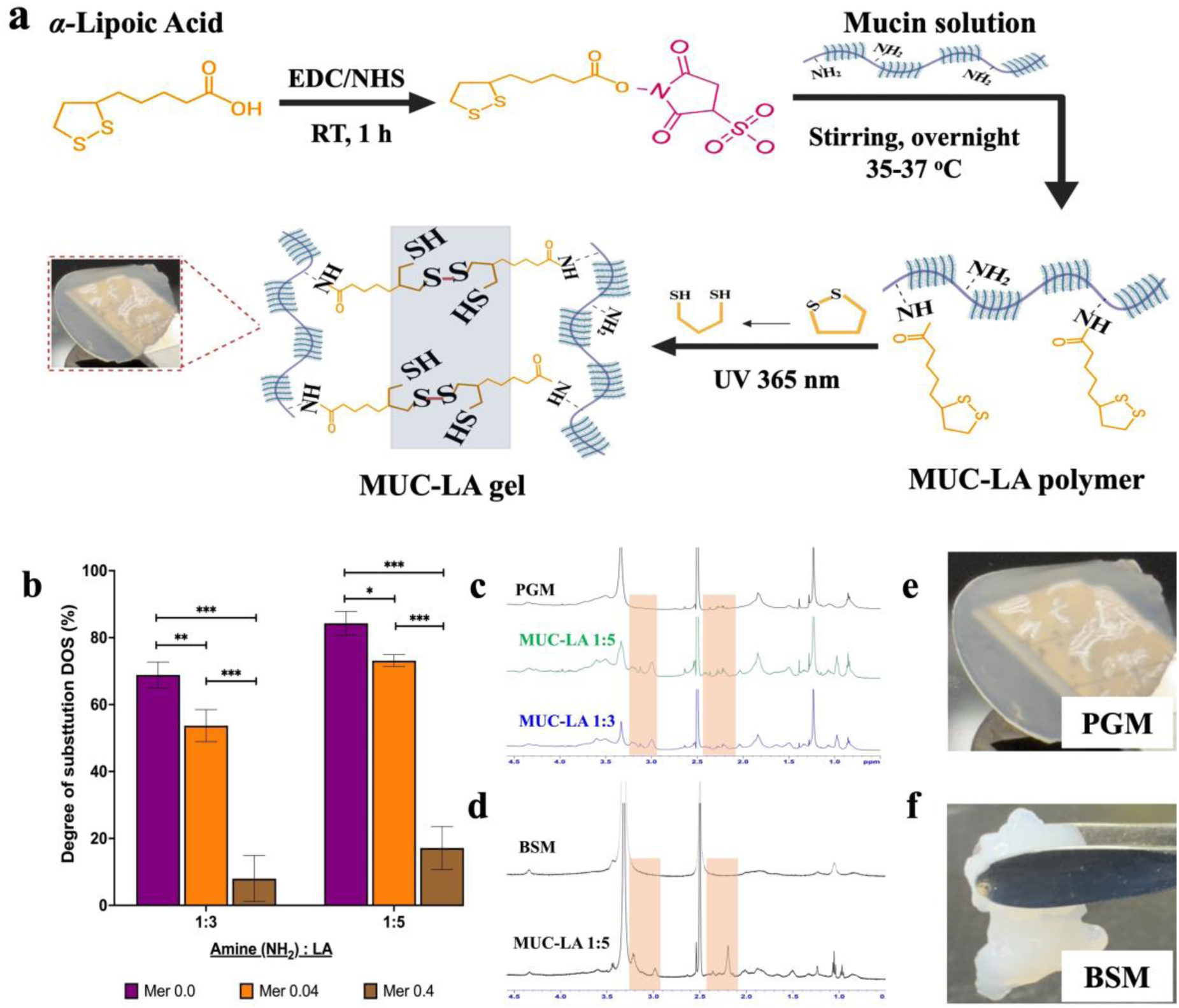
Synthesis and characterization of lipoic acid (LA) modified mucin. (a) Schematic representation of the conjugation of LA onto mucin backbone chains to synthesize mucin-LA (MUC-LA) polymer. Resultant MUC-LA polymer undergoes dithiolane-mediated photo-crosslinking under UV 365 nm light, forming a disulfide crosslinked hydrogel. (b) Degree of substitution DOS (%) of LA onto porcine gastric mucin (PGM) as a function of amine-to-LA molar ratio and various mercaptoethanol (Mer) concentrations. Reactions were carried out at two amines:LA molar ratios (1:3 and 1:5) in the presence of different EDC-Mer molar ratios (0.0, 0.04, and 0.4). The DOS (%) was quantified by colorimetric accessible amine group analysis after conjugation. Data are presented as mean ± SD (*n* = 3), using one-way ANOVA, Tukey’s multiple comparison test (\**p* < 0.05, \*\**p* < 0.01, \*\*\**p* < 0.001). (c) ¹H NMR spectra of unmodified PGM and modified PGM/MUC-LA synthesized using amine:LA feed ratios of 1:3 and 1:5. (d) ¹H NMR spectra of unmodified bovine submaxillary mucin (BSM) and modified BSM/MUC-LA using amine:LA feed ratio of 1:5. Highlighted regions indicate resonances assigned to the dithiolane ring and aliphatic protons of conjugated LA. (e, f) Representative photographs of the photo-crosslinked 5% (w/v) PGM-LA/BSM-LA hydrogel following UV irradiation, respectively.

Accessible thiol (-SH) and primary amine (-NH_2_) groups were first quantified in commercial and re-purified porcine gastric mucin (PGM) to identify suitable sites for subsequent functionalization **(Figure S1a,b, Supporting Information)**. Re-purification reduced the accessible thiol content, whereas a substantial fraction of primary amines remained available, supporting the use of primary amines as the preferred conjugation sites for LA. To evaluate the extent of LA conjugation to the mucin backbone, the degree of substitution (DOS) was calculated by quantifying the decrease in accessible primary amine groups using the TNBSA assay. As shown in **Figure 2b**, increasing the amine:LA molar ratio from 1:3 to 1:5 increased the DOS from ≈68.85% to ≈84.28%, respectively, indicating that higher the LA availability promoted more extensive functionalization of mucin. This trend is consistent with previous studies on gelatin and polysaccharide-based systems, where increasing LA availability led to higher levels of backbone functionalization [29–30]. We also investigated the effect of *β*-mercaptoethanol (Mer), which has been used previously to quench the residual EDC in carbodiimide coupling reactions [29]. In contrast to the reported studies in simpler proteins, the highest DOS (%) values with *no* Mer added for both amine:LA ratios were obtained in our system [29]. Conversely, the addition of 0.04 or 0.4 equivalents of Mer relative to EDC reduced the DOS to ≈53.71% and ≈8% for 1:3, and to ≈73.14% and ≈17.14% for 1:5, respectively. These results suggest that Mer interfered with LA coupling under the conditions used. The reduced DOS (%) in the presence of Mer may also reflect the limited accessibility of reactive amines within the highly glycosylated and heterogenous mucin backbone, suggesting that coupling conditions optimized for simpler proteins may not apply directly to the mucins and may require further optimization [31–32]. Chemical conjugation of LA to the mucin backbone was further assessed by ¹H NMR spectroscopy (500 MHz, Avance III HD, Bruker, Germany). Compared with unmodified mucins PGM and bovine submaxillary mucin (BSM), the modified MUC-LA samples exhibited additional or enhanced resonance in the aliphatic regions as indicated by the highlighted regions **(Figure 2c,d)**. These signals were observed under both (1:3 and 1:5) conjugation conditions, supporting the reproducible functionalization of the mucin chains. These additional resonances are consistent with the characteristic aliphatic signals of LA-derived structures reported previously [33]. Following UV irradiation at 365 nm, both 5% (w/v) PGM-LA and BSM-LA precursors formed hydrogels as shown in the representative photographs in **Figure 2e,f**. Together, these results establish that LA can be reproducibly introduced onto both PGM and BSM and that the extent of functionalization can be tuned through the reaction conditions. The introduced dithiolane groups provide the chemical basis for the photo-induced network formation and redox-responsive behavior examined in the following sections.

### 2.2 Photo-Crosslinking of Lipoic Acid Modified Mucin Hydrogels and Bulk Rheological Characterization

The photo-gelation of PGM/mucin-LA (MUC-LA) hydrogels was monitored in real time using a UV-integrated rheometer (MCR 302WESP, Anton Paar, Austria) **(Figure 3a)**. Before irradiation, both storage and loss modulus, (G′) and (G″) respectively, remained low, reflecting the sol-like state of the precursor solutions. Exposure to UV light (365 nm, 20 mW cm^-2^) caused a rapid increase in both moduli, G′ crossing G″ within approximately 10-50 s and reached plateau within 5-10 mins depending on the concentrations such as 1.25%, 2.5%, 3% and 5% (w/v) **(Figure 3b,c)**. This rapid sol-to-gel transition is attributed to photo-induced opening of the 1,2-dithiolane groups followed by intermolecular disulfide formation between mucin chains without adding any photo-initiator [30]. Higher MUC-LA concentrations produced larger plateau G′ values, indicating concentration-dependent control over network strength. **Supplementary Video S1** shows the macroscopic flow behavior of the 3% (w/v) MUC-LA hydrogel under applied deformation, consistent with its soft viscoelastic response.

**Figure 3.**
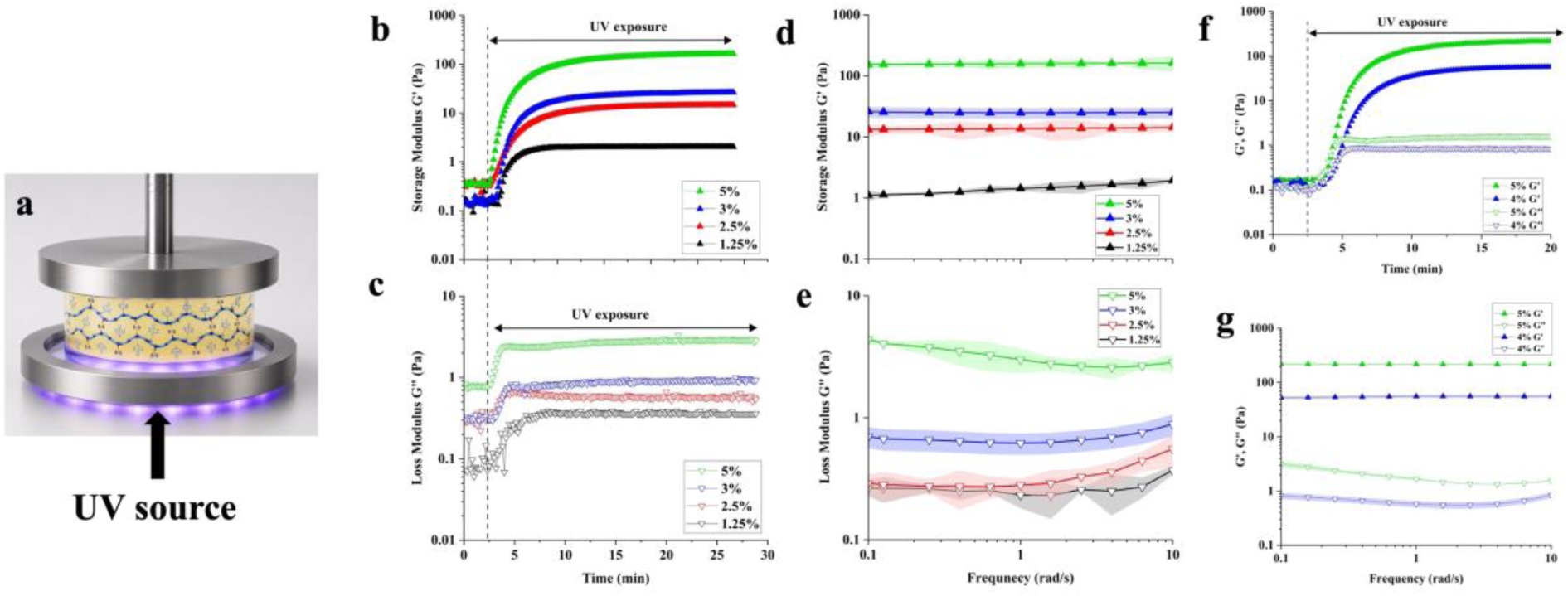
Rheological characterization of PGM-LA and BSM-LA hydrogels. (a) Schematic illustration of the UV-integrated rheometer configuration used for in situ photo-gelation. (b-c) Rheological characterization of PGM-LA hydrogels at different mucin concentrations, including time-sweep measurements of G′ (filled triangles) and G″ (unfilled triangles) during UV irradiation and the corresponding (d,e) frequency-sweep G′ and G″ after complete gel formation. (f-g) Rheological characterization of BSM-LA hydrogels at different mucin concentrations, showing UV-induced evolution of G′ and G″ during gelation and the frequency-dependent G′ and G″ of the resulting networks. Data are presented as mean ± SD (*n* = 3) and time-sweep curves are shown as representative traces without error bands for clarity.

The mechanical properties of resulting hydrogels were further examined by frequency-sweep measurements **(Figure 3d,e)**. Across 0.1-10 rad s⁻¹, G′ remained consistently higher than G″ and showed only weak frequency dependence, indicating a predominantly elastic network. At low frequency (0.1 rad s^-1^), G′ values increased from ≈1-2 Pa for the 1.25%, ≈ 13.14 Pa for 2.5%, ≈26.06 Pa for 3% and ≈153 Pa for the 5% hydrogel formulations, showing a clear concentration-dependent increase in storage modulus.

The concentration-dependent increase in modulus is consistent with the general behavior of mucin-based networks, in which intermolecular associations and entanglement are mostly contributed by disulfide crosslinking to bulk rheological properties [25, 34]. The modulus range of the PGM-LA formulations falls within mechanically relevant regimes reported for biological mucus. Healthy mucus generally exhibits lower elastic moduli, whereas hyper-concentrated mucus in conditions such as cystic fibrosis becomes stiffer and more restrictive to transport [35]. Similar disease-associated changes in mucus viscoelasticity have also been reported in the intestine [36]. These findings indicate that PGM-LA can reproduce a broad range of mucus-relevant viscoelastic properties while maintaining a predominantly elastic response.

To assess whether this approach could be extended beyond PGM, BSM was also functionalized with LA and evaluated under the same UV-triggered conditions **(Figure 3f)**. BSM-LA similarly underwent rapid gelation and became elastic (G′ > G″) during frequency-sweep analysis. The G′ value at 0.1 rad s^-1^ observed as ≈52.74 Pa and ≈219.17 Pa for 4% (w/v) and 5% (w/v) hydrogel concentration, respectively **(Figure 3g)**. Although BSM is derived from a different biological source than PGM [37], its successful conversion into a hydrogel supports the transferability of the LA-mediated photo-gelation strategy across distinct mucin types. Together, these results establish LA functionalization as a versatile route for generating rapidly photo-crosslinked and mechanically tunable mucus-mimetic materials from commercially available mucins.

### 2.3 Redox-Responsive and Self-Healing ability of MUC-LA Hydrogels

The disulfide-crosslinked PGM/MUC-LA network is expected to respond to a reducing environment as illustrated in **Figure 4a**. To evaluate this response, 4% (w/v) prepared hydrogels were exposed to glutathione (GSH) for 24 h at 10 and 20 mM for a clear difference, while PBS-treated samples served as the control and measured at 0 and 24 h. Frequency-sweep measurements showed a clear reduction in G′ after GSH treatment, with values of ≈17.42 Pa and ≈14.53 Pa for 10 and 20 mM, respectively, compared with the control showing ≈45.16 Pa and 29.12 Pa at 0 and 24 h, respectively **(Figure 4b)**. This decrease in G′ indicates network softening under reducing conditions and is consistent with reduction and/or exchange of disulfide junctions, which can reduce effective network connectivity. Such redox sensitivity is relevant to mucus, where disulfide bonds contribute to mucin-network mechanics and can be altered by reducing agents [38–39].

**Figure 4.**
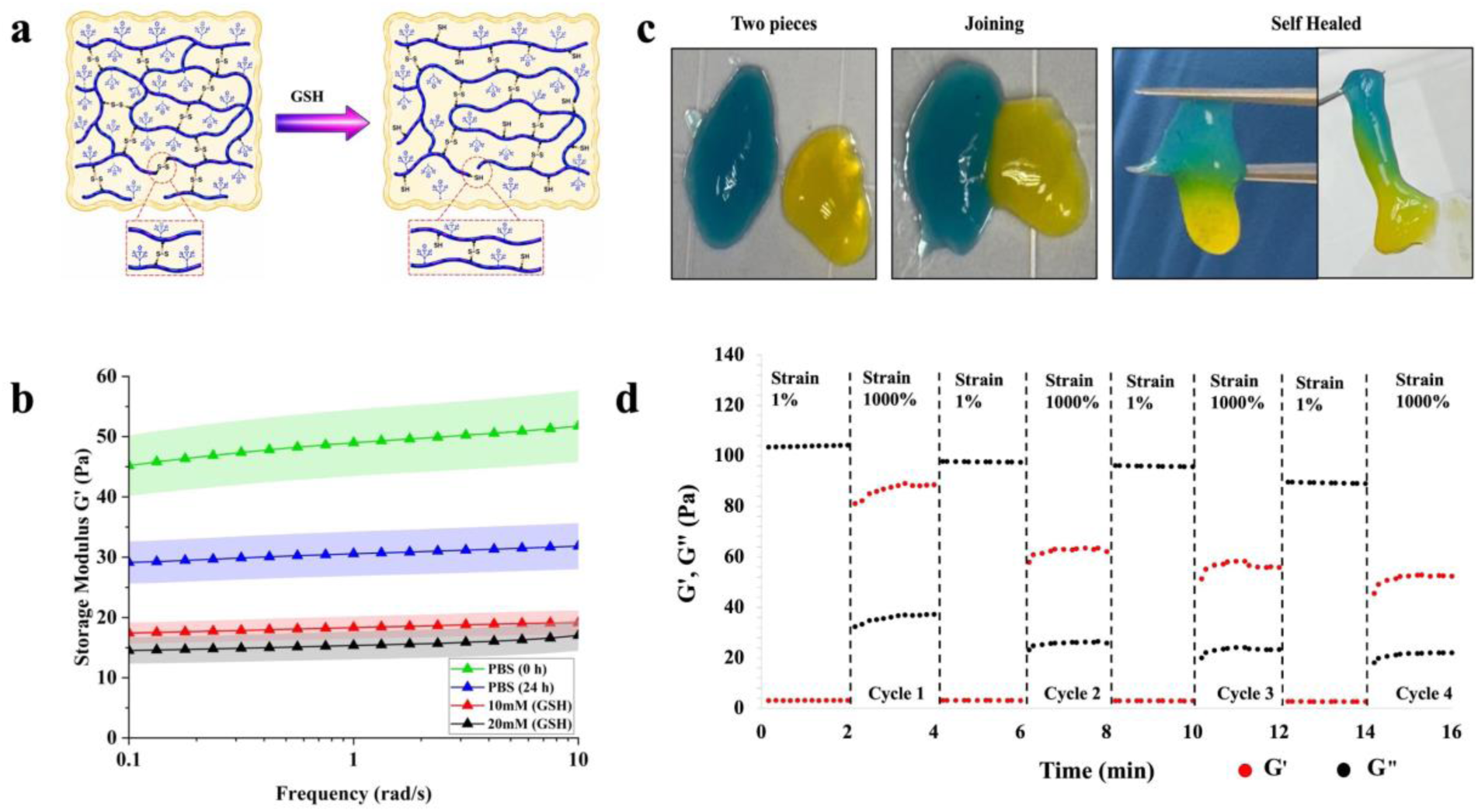
Redox-responsive and self-healing behavior of PGM/MUC-LA hydrogels. (a) Schematic representation of GSH-mediated reduction of disulfide crosslinks within the MUC-LA network. (b) Frequency-dependent G′ of hydrogels following exposure to PBS or GSH (10 and 20 mM), showing network softening under reducing conditions. (c) Macroscopic self-healing of two differently colored hydrogel segments following direct interfacial contact. (d) Cyclic step-strain rheology of 5% (w/v) hydrogel under alternating low (1%) and high (1000%) strain over four consecutive cycles, demonstrating reversible network disruption and substantial recovery of the elastic response. Black and red dots represent G′ and G″, respectively.

Self-healing behavior was next evaluated for MUC-LA hydrogel. Two separately prepared hydrogel pieces containing different color additives were placed in direct contact **(Figure 4c)**. After 1-2 h at room temperature, the interface became continuous and the two pieces could be lifted together, indicating recovery of cohesion across the contact surface. Mechanical recovery was further quantified by cyclic step-strain rheology **(Figure 4d)**. A prior amplitude sweep showed network yielding above approximately 500% strain **(Figure S2, Supporting Information)**, therefore, 1000% strain was used to disrupt the 5% (w/v) hydrogel network and 1% strain to monitor recovery. At 1% strain, G′ was approximately 100 Pa. Increasing the strain to 1000% caused G′ to drop below G″, indicating the loss of the predominantly elastic response. When the strain was returned to 1%, G′ rapidly recovered to approximately 80-90% of its initial value. This recovery was maintained over four consecutive cycles, demonstrating repeated restoration of the network after large deformation. Complete recovery was not achieved, given the limited recovery time of ∼2 min. The combined GSH-induced softening, macroscopic interfacial healing and cyclic rheological recovery show that MUC-LA exhibits both chemical responsiveness and mechanical reversibility. This behavior is consistent with dynamic disulfide exchange, which can support network rearrangement after chemical or mechanical disruption [27]. These properties are particularly relevant for mucus-mimetic materials, where reversible interactions help preserve network function under changing chemical and mechanical conditions [23, 40].

### 2.4 Size-Dependent Nanoparticles Transport Restriction within MUC-LA Hydrogels

To probe the microscale transport within the PGM/MUC-LA hydrogel, PEG-coated nanoparticles of different sizes were used as tracer particles. This approach enabled assessment of how particle size influences mobility through the hydrogel matrix. PEGylation was confirmed by small increases in hydrodynamic diameter and clear changes in surface charge. Following PEGylation, the nominal diameter of 100 and 200 nm particles increased from 118.4 ± 1.75 to 138.1 ± 4.2 nm (mean ± SD) and from 204.5 ± 3.87 to 225.9 ± 1.05 nm, respectively, determined by Zetasizer Ultra (Malvern Instruments, UK), and the corresponding zeta potentials changed from –56.32 ± 0.34 to –4.14 ± 2.12 mV (mean ± SD) and from –59.41 ± 2.61 to –1.88 ± 3.10 mV, respectively, indicating the charge neutrality of particles after the modification. These changes are consistent with PEG surface coating without major particle aggregation and agree with previous reports using PEGylated polystyrene probes used in mucus transport studies [41]. PEGylation was used to reduce nonspecific adhesive interactions with mucin and thereby improve the assessment of particle transport within the hydrogel network. The resulting particles were then tracked within a 3% (w/v) MUC-LA hydrogel to examine microscale transport through the network **(Figure 5a)**. Particle trajectories were extracted using a subpixel localization and tracking workflow, with representative localization analysis shown in **Figures S3 and S4, Supporting Information**. Representative trajectories showed a clear difference in particle mobility. The 100 nm particles explored substantially larger regions, whereas the 200 nm particles remained highly localized **(Figure 5b,c)**. Representative side-by-side recordings further illustrate the greater mobility of the 100 nm particles and the more restricted motion of the 200 nm particles within the 3% (w/v) MUC- LA hydrogel **(Supplementary Video S2)**. This difference was quantified by mean-squared displacement (MSD) analysis. MSD analysis and power-law fitting were restricted to lag times (*τ*) ≤1s to minimize statistical uncertainty at longer lag time *τ*. Individual trajectories of the 100 nm particles (*n* = 31) showed substantially larger MSD values that increased with lag time *τ*, whereas the 200 nm particles (*n* = 53) showed much smaller displacements with limited time dependence **(Figure 5d)**.

**Figure 5.**
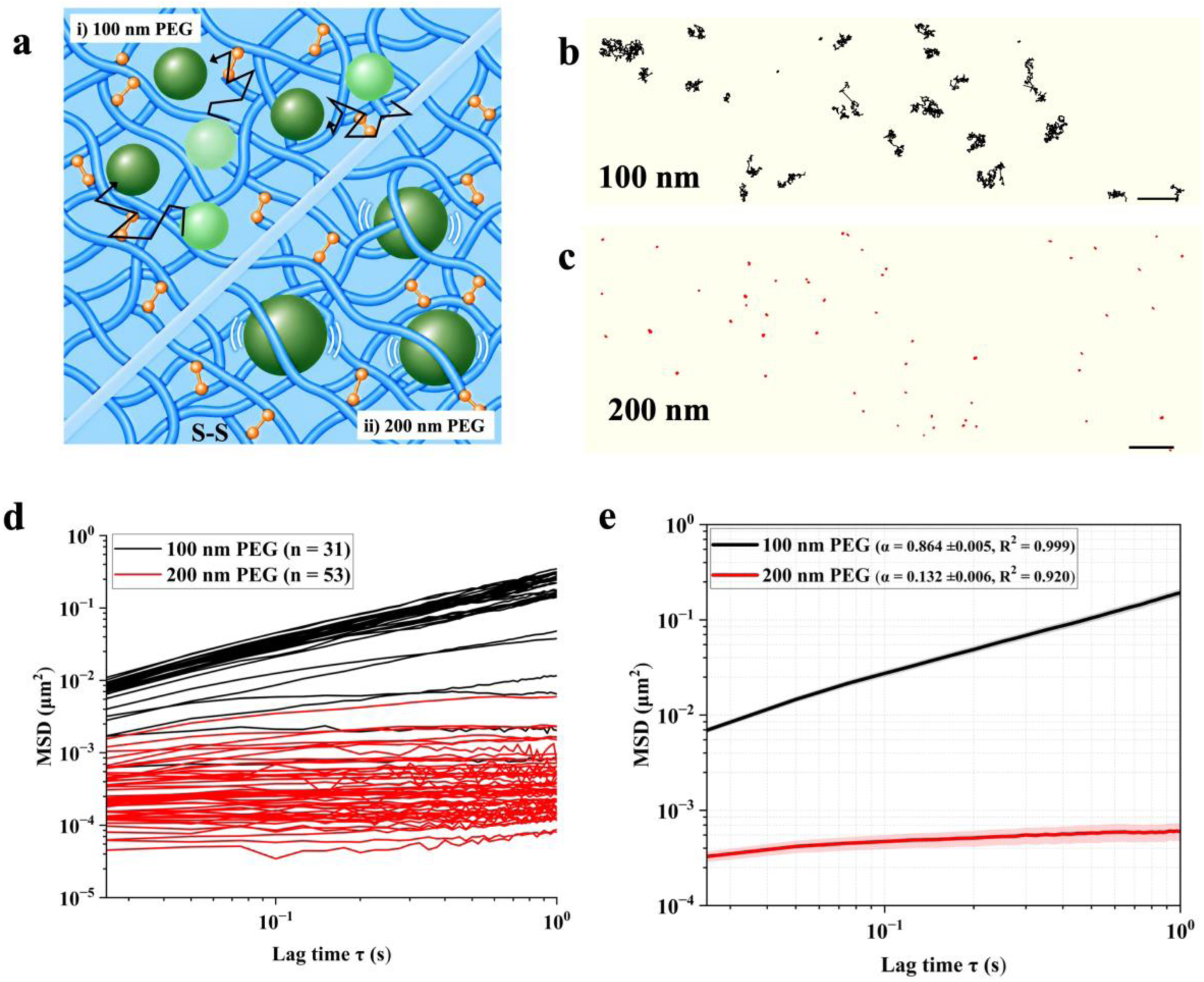
Size-dependent nanoparticles movement within the MUC-LA hydrogel network. (a) Schematic illustration of PEG-coated fluorescent nanoparticles with diameters of 100 and 200 nm within the disulfide-crosslinked mucin network. (b,c) Representative trajectories of both PEG-coated nanoparticles over 60 s recorded videos. (d) Individual mean-square displacement (MSD) profiles of 100 nm (*n* = 31) and 200 nm (*n* = 53) particles as a function of lag time *τ*. (e) Ensemble-averaged MSD profiles fitted using the power-law relation *MSD* ∝ *τ^α^*, with the corresponding anomalous diffusion exponents and R^2^ values indicated in the panel.

Ensemble MSD analysis further highlighted this difference, as shown in **Figure 5e**. Power-law fitting yielded an anomalous diffusion exponent of *α* = 0.864 ± 0.005 (R^2^ = 0.999) for the 100 nm particles, indicating relatively mobile but sub-diffusive motion. In contrast, the 200 nm particles exhibited a substantially lower *α* = 0.132 ± 0.006 (R^2^ = 0.920), indicating strongly restricted transport within the hydrogel.

These results show that particle mobility is strongly size-dependent, suggesting that the MUC-LA network acts as a size-selective barrier, with the nominal range of mesh size (*ξ*) around 100-200 nm. Comparable behavior has been reported in purified mucin and native mucus systems, where nanoparticle transport is influenced by particle size together with steric and physicochemical interactions with the mucin network [41–43]. Together with the bulk rheological measurements, these findings show that MUC-LA exhibits mucus-relevant viscoelastic behavior (G′ > G″) and mesh size within the native mucus range reported earlier [44]. The pronounced restriction of the larger nanoparticles further motivated us to evaluate the MUC-LA network as a barrier to viral transport.

### 2.5 Barrier Properties of MUC-LA Hydrogels against SARS-CoV-2 and Influenza A Virus Viral Infections

Given that the mucin hydrogel rectifies the translocation of 100 and 200 nm PEG-coated particles shown in **Figure 5**, we next examined whether this physical barrier could be leveraged to block virus transmission. Prior to the viral studies, we first checked the cytocompatibility of both gel- and solution of PGM/MUC-LA, and the results showed no evident reduction in MDCK cell viability under the tested conditions, supporting the cytocompatibility of the material **(Figure S5, Supporting Information)**. To this end, we established a transwell-based infection assay to evaluate the efficacy of mucin hydrogel in preventing viral infection **(Figure 6a)**. The hydrogel, alongside control solutions of PBS and non-crosslinked mucin, was layered onto the membrane of transwell, which was inserted into a 24-well plate seeded with Vero E6 cells. A viral inoculum containing SARS-CoV-2 was then gently added atop these materials, carefully avoiding premature mixing, to test whether the virus could diffuse through the layer to infect the underlying cells. Vero E6 cells infection was assessed by immunofluorescence assay through the recognition of viral nucleocapsid (N) protein **(Figure 6b)**.

**Figure 6.**
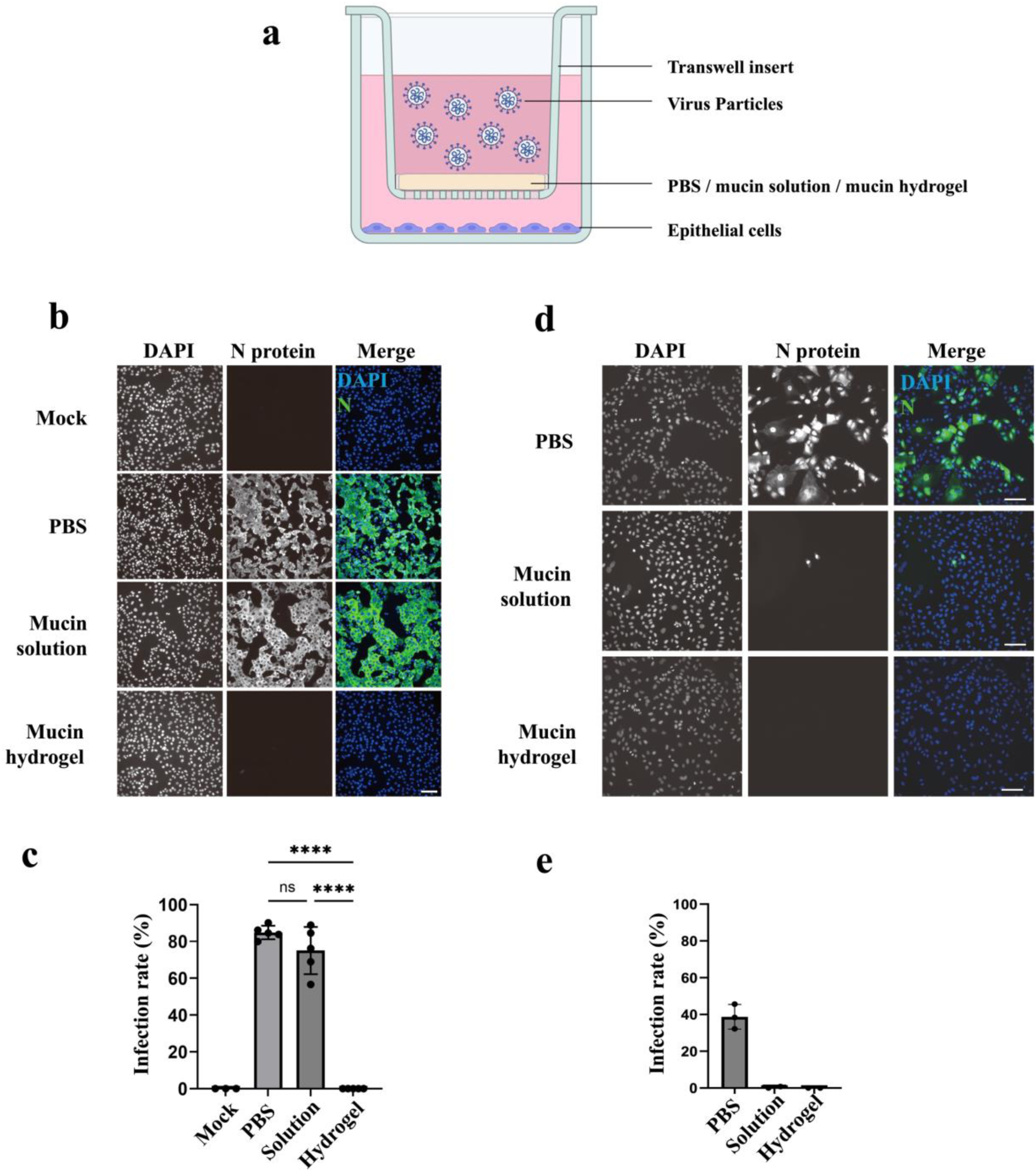
PGM/MUC-LA hydrogel (3% w/v) restricts transmission of SARS-CoV-2 and influenza A virus (IAV) across a transwell barrier. (a) Schematic of the transwell-based viral barrier assay, with virus particles introduced into the apical compartment above PBS, mucin/MUC-LA solution, or in situ MUC-LA hydrogel and target epithelial cells cultured in the basolateral compartment. (b) Representative immunofluorescence images of SARS-CoV-2 infection in Vero E6 cells at 24 h post-inoculation. (c) Quantification of SARS-CoV-2 infection under the indicated barrier conditions. (d) Representative immunofluorescence images of IAV infection in MDCK cells at 16 h post-inoculation. (e) Quantification of IAV infection under the corresponding conditions. Viral N/NP protein is shown in green, and nuclei are counterstained with DAPI (blue); scale bars, 100 µm. Each data point represents an independent biological replicate, calculated from 2-3 randomly selected fields of view for SARS-CoV-2 and 1-2 fields for IAV. Statistical significance was determined by one-way ANOVA followed by Tukey’s multiple-comparisons test; \*\*\*\**p* ≤ 0.0001; ns, not significant.

At 24 hours post-inoculation, infection rates were high in both the PBS control (84.90 ± 3.67%, mean ± SD) and the non-crosslinked 3% (w/v) MUC-LA control (75.05 ± 12.82%, mean ± SD), with no statistically significant difference between them. In contrast, no SARS-CoV-2-infected cells were detected in the presence of the 3% (w/v) MUC-LA hydrogel (**Figure 6c**). Protection also depended on MUC-LA concentration: the 2% (w/v) hydrogel yielded a low but detectable infection rate of approximately 2.27%, whereas no infection was detected with the 3% formulation (**Figure S6, Supporting Information**). This concentration dependence is consistent with greater restriction of viral transport at higher MUC-LA concentrations, although the contribution of changes in network structure requires further investigation. Together, these results support an important role for network formation in protecting the underlying epithelial cells against SARS- CoV-2 infection under the tested conditions.

To assess whether protection extends to another virus, we examined influenza A virus (IAV). Both the 3% (w/v) MUC-LA hydrogel and non-crosslinked MUC-LA strongly suppressed IAV infection (**Figure 6d,e**), suggesting that protection against IAV does not require crosslinking. IAV attachment to host cells involves binding of viral hemagglutinin (HA) protein to the sialic acid receptor on cell surface [45]. Sialic acid residues on mucin [46] may therefore act as decoy binding sites for HA, reducing viral attachment to epithelial cells. This proposed mechanism remains to be established directly but is consistent with previous reports that mucus and mucus-inspired hydrogels restrict viral mobility through physical and biochemical interactions [20, 47].

Overall, these findings demonstrate that crosslinked MUC-LA protects epithelial cells against both SARS-CoV-2 and IAV in transwell-based infection models. The contrasting effects of crosslinked and non-crosslinked MUC-LA suggest virus-dependent contributions from network formation and mucin-mediated interactions. These results support MUC-LA hydrogels as mucus-mimetic protective barriers and motivate further investigation of their antiviral mechanisms and activity against additional viruses.

## 3 Conclusions

Commercially available mucins were directly functionalized with α-lipoic acid to introduce photoresponsive dithiolane groups, enabling the formation of single-component, disulfide-crosslinked hydrogels under 365 nm UV irradiation without added any photo-initiators or external polymeric crosslinkers. Inspired by the disulfide-linked architecture of native mucins, this strategy enabled the reconstruction of mucus-mimetic MUC-LA hydrogels with tunable viscoelasticity, self-healing behavior, and glutathione-responsive softening. Tracking of PEGylated nanoparticles revealed greater mobility of 100 nm than 200 nm, demonstrating size-dependent restricted transport within the hydrogel network. Importantly, in transwell-based infection models, no SARS-CoV-2-infected epithelial cells were detected beneath the soft 3% (w/v) MUC-LA hydrogel, whereas infection remained high with the non-crosslinked precursor, reflecting the protective role of native mucus. These findings support an important contribution of network formation to SARS-CoV-2 protection. In contrast, both non-crosslinked and crosslinked MUC-LA suppressed influenza A virus infection, suggesting that mucin-mediated interactions may contribute to protection independently of crosslinking. Together, these results establish MUC-LA as an experimentally controllable mucus-mimetic platform that integrates tunable mechanics, dynamic disulfide chemistry, size-dependent transport and protection against viral infection. The current platform offers opportunities to investigate mucosal transport, host-pathogen interactions and therapeutic delivery across mucus-like barriers. These dynamic properties may further inspire tissue engineering and injectable hydrogel applications, subject to further evaluation of cytocompatibility, injectability and stability under physiological conditions.

## Experimental Section

### Synthesis and Characterization of Lipoic Acid Conjugated Mucin

LA was covalently conjugated to accessible amine groups on mucin backbone using EDC/NHS carbodiimide chemistry, adapted with necessary modifications from a reported gelatin-based protocol [29]. Re-purified PGM (0.5 g) was first dispersed in DMSO and then diluted with 0.1 mol L^-1^ bicarbonate buffer (pH 8-9) to mucin concentration of approximately 12 mg mL⁻¹. Separately, LA was dissolved in DMSO at 30 mg mL⁻¹, while EDC/NHS dissolved in 0.1 mol L^-1^ MES buffer (pH 6.5). Later, EDC/NHS were added to the LA solution at a fixed LA:EDC:NHS molar ratio of 1:3:3 and stirred at 250 rpm, 1 h at room temperature (RT) for activating LA carboxyl groups. Where indicated, *β*-mercaptoethanol (Mer) was added at 0, 0.04 and 0.4 molar ratios to EDC. The activated LA mixture was subsequently added to the mucin solution under continuous stirring, giving a final mucin concentration of approximately 10 mg mL⁻¹, DMSO 10-12% (v/v), and LA ≤ 3 mg mL⁻¹ to minimize apparent precipitation. The reaction mixture was incubated at 37 °C overnight, with the pH maintained between 8-9 during the first 2-3 h to favor amide bond formation.

Subsequently, the solution mixture was centrifuged at 1000 rpm around 10-15 min to remove insoluble material and mucin aggregates. The supernatant was purified by dialysis using a 12-14 kDa MWCO membrane for 3 days, with the regular exchange of solvent three times a day. The purified MUC-LA solution was freeze-dried and stored at -20 °C. The degree of substitution DOS (%) was calculated from the decrease in accessible primary amine (-NH_2_) groups measured by the TNBSA assay relative to unmodified mucin. To evaluate the tunability of LA substitution, mucin amine:LA molar ratios of 1:3 and 1:5 were investigated while keeping the LA:EDC:NHS ratio constant at 1:3:3. Successful LA conjugation was further assessed by ^1^H NMR spectroscopy (500 MHz, Avance III HD, Bruker, Germany) using D₂O as the solvent for unmodified PGM and modified PGM/MUC-LA samples prepared at amine:LA ratios of 1:3 and 1:5. The same conjugation procedure was applied to BSM using an amine:LA ratio of 1:5. For BSM, LA conjugation was evaluated by only ^1^H NMR spectroscopy.

### Bulk Rheological Characterization of UV-Induced Gelation

The rheological properties of MUC-LA hydrogel were determined using a rheometer (MCR 302WESP, Anton Paar, Austria) equipped with a custom integrated UV light source (Brightek, Taiwan), enabling real-time monitored photo-induced gelation. Measurements were performed using a 25 mm parallel-plate (pp-25) geometry at 25 °C. A serrated parallel plate geometry was used to minimize wall slip, particularly during large amplitude oscillatory shear and self-healing measurements. The MUC-LA precursor solution in PBS was loaded between the plates at a 1 mm gap. Samples were equilibrated for 2-3 min, followed by UV light (365 nm, 20 mW cm^-2^) to initiate photo-gelation. The evolution of the storage (G′) and loss (G′′) modulus was monitored in real time, keeping a strain of 1% and angular frequency of 10 rad s⁻¹ until the moduli reached a plateau. Oscillatory strain sweep analysis was used for the determination of hydrogels linear viscoelastic region (LVER). The G′ and G″ values of hydrogels were determined using frequency-sweep (0.1-10 rad s⁻¹) at 1% strain.

### Redox Responsive and Self-Healing Properties

The MUC-LA hydrogel contains dynamic disulfide bonds that are responsive to reducing conditions such as glutathione (GSH). Redox responsiveness was evaluated by incubating fully formed PGM/MUC-LA 4% (w/v) hydrogels with GSH at 10 and 20 mM for 24 h at RT, while untreated hydrogels maintained in PBS under identical conditions served as controls. Following incubation, changes in the mechanical properties of hydrogels were evaluated using frequency-sweep measurements.

An oscillatory strain sweep was first performed to determine the onset of network yielding before cyclic self-healing measurements. Mechanical recovery was then evaluated by cyclic step-strain rheology. The hydrogel was subjected to alternating low and high strain cycles 1% and 1000% respectively, at constant frequency 10 rad s^-1^ to evaluate network disruption and recovery. Each cycle consisted of 2 min of 1% followed by 2 min of 1000% strain, and the cycles were repeated four times. In addition to rheological evaluation, self-healing ability of the MUC-LA hydrogel was visually assessed using colored hydrogel segments. Two separate MUC-LA hydrogel samples were prepared by incorporating fluorescein sodium salt and methylene blue to generate green and blue hydrogel segments, respectively. The two segments were then brought into direct contact and incubated at RT for 1-2 h before visual assessment of interfacial healing.

### Preparation and Characterization of PEGylated Polystyrene Nanoparticles

Yellow-green fluorescent carboxylate-modified polystyrene microspheres with nominal diameters of 100 and 200 nm (excitation/emission: 505/515 nm) were PEGylated by EDC/NHS-mediated coupling of 5 kDa methoxy-poly(ethylene glycol)-amine (mPEG-NH₂), following a previously reported procedure [41]. Reactions were carried out in HEPES-buffered saline (HBS; 10 mM HEPES, 150 mM NaCl, pH 8.0) containing 0.7% particle solids, 10 mg mL⁻¹ mPEG-NH₂, 4 mg mL⁻¹ EDC, and 1.13 mg mL⁻¹ NHS. Samples were protected from light and incubated overnight at RT, while shaking at 200 rpm. Excess reagents were removed using 100 kDa Amicon Ultra centrifugal filters (Millipore, MA, USA) by centrifugation at 14,000 rpm for 12 min, followed by two HBS washes. PEGylated particles were recovered by reverse centrifugation at 1,000 rpm for 3 min and resuspended in HBS. Hydrodynamic diameter and zeta potential were measured before and after PEGylation using a Zetasizer Ultra (Malvern Instruments, UK).

### Particle Tracking Analysis

PEGylated 100 and 200 nm particles were dispersed separately in 3% (w/v) MUC-LA precursor at final concentrations of 1×10^8^ particles mL^-1^. Approximately 20-30 µL of each sample was loaded into individual channels of µ-Slide VI 0.4 (Ibidi, Germany). The reservoirs were sealed to minimize evaporation, and the samples were UV-crosslinked under the same conditions used for bulk rheology (365 nm, 20 mW cm⁻²). After equilibration for approximately 30 min, particles movement was recorded at RT using a Leica DMI 4000 B microscope equipped with a 100x oil-immersion objective and an Andor iXon DV887_BV EMCCD camera. Videos were acquired using frame transfer mode at 40 frames s^-1^ with an exposure of 24.52 ms and a calibrated image scale of 0.160 µm pixel⁻¹. For each particle size, 4-5 regions were recorded approximately 40-50 µm above the channel surface.

Particle candidates were detected and initially linked using TrackPy following established digital particle-tracking approaches [48]. Final subpixel coordinates were obtained using a custom Python routine by fitting the fluorescence intensity within a recentered 7 × 7 pixel region of interest (ROI) to a centrosymmetric two-dimensional Gaussian model **(Figure S3, Supporting Information)** [49–50]. The fluorescence intensity distribution was described by Equation (1):

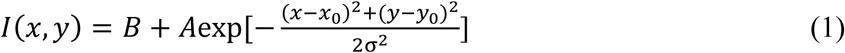

where *B* is the background intensity, *A* is the fitted amplitude, *x_0_* and *y_0_* are the fitted particle centroid coordinates and *σ* represents the Gaussian width.

Localization uncertainty was estimated from the covariance matrix of the Gaussian fit **(Figure S4, Supporting Information)**. For each accepted trajectory, the two-dimensional time-averaged mean-squared displacement (MSD) was calculated from particle displacements at defined lag times [51–52]. The measured MSD was calculated using Equation (2):

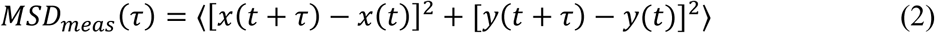

where τ is the lag time and the brackets denote the averaging over all displacement pairs available at that lag time.

To account for localization uncertainty, the positional uncertainties associated with both endpoints of each displacement were propagated as described previously [52–53]. The propagated localization variance was calculated using Equation (3):

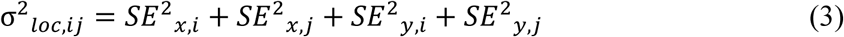

where 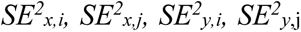 are the localized standard errors for the two particle positions (x and y) defining each displacement.

The corrected MSD was then obtained using Equation (4) by subtracting the mean propagated localization variance at each lag time from the measured MSD:

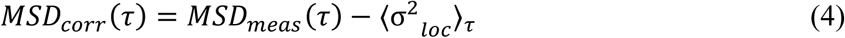

Individual-particle MSD curves were averaged with equal particle weighting and presented as mean ± SEM. After applying the trajectory quality-control (QC) criteria described in the Section 2.4, **Supporting Information**, 31 and 53 trajectories were retained for 100 and 200 nm particles, respectively.

Anomalous transport was characterized using the power-law relationship in Equation (5):

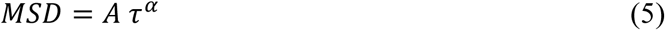

where *A* is the generalized transport coefficient and α is the anomalous diffusion exponent.

Power law fitting was performed over lag time τ = 0.025-1.0 s to reduce statistical uncertainty associated with the decreasing number of displacement pairs at longer lag times.

### Barrier Properties Against Virus

#### Cell Culture

In this study, African green monkey kidney Vero E6 cells (CRL-1586, ATCC) were used for SARS-CoV-2 (Wuhan strain) infection, and Madin-Darby canine kidney (MDCK) cells (60004, BCRC, Taiwan) were used for influenza A virus (IAV, A/Puerto Rico/8/1934(H1N1) strain) infection. Both cell lines were cultured in MEM supplemented with 10% FBS and incubated at 37°C in a humidified atmosphere with 5% CO_2_.

#### Transwell-Based Infection Assay

Vero E6 and MDCK cells were seeded onto 12 mm glass coverslips in 24-well plates at 1.1 x 10^5^ or 1.0 x 10^5^ cells/well, respectively. The cells were cultured for 18 hours at 37 °C with 5% CO_2_ to allow adequate attachment and achieve the desired confluency. On the day of infection, cells were washed once with 1 mL of PBS, and the culture medium was replaced with 700 μL of MEM supplemented with 0.3% BSA and 1× P/S. Transwells (SPLInsert™ Hanging, SPL Life Sciences, Pocheon, Korea) were loaded with PBS, mucin solution, or mucin hydrogel (35 μL for SARS-CoV-2 assays, 1 mm gel thickness; 50 μL for IAV assays, 1.5 mm gel thickness), and then installed into 24-well plates. Subsequently, viruses were added into the apical chamber of the transwell in a total volume of 150 μL. In SARS-CoV-2 infection experiments, 1.1 x 10^6^ plaque-forming units (PFU) of virus was used, corresponding to an estimated multiplicity of infection (MOI) of 10 based on unrestricted viral passage through the transwell. In the case of IAV infection, 5.0 x 10^5^ PFU of virus was used, corresponding to MOI of 5 upon unrestricted viral inoculation. The whole transwell-based infection system was incubated for 24 hours following SARS-CoV-2 infection or 16 hours following IAV infection. At the indicated endpoints of viral infections, the transwells were removed, and the cells in the lower chamber were fixed with 4% PFA for 10 minutes at room temperature. Subsequently, the cells were subjected to immunofluorescence assay to assess the infection rate. While most conditions were independently repeated, the 2% (w/v) mucin hydrogel/SARS-CoV-2 infection group was tested only once.

#### Immunofluorescence Assay

PFA-fixed cells were quenched with 100 mM glycine in PBS for 5 minutes, permeabilized with 0.1% Triton X-100 in PBS for 10 minutes and blocked with 5% goat serum in PBS for 1 hour to prevent non-specific binding. For viral protein detection, cells were incubated overnight at 4 °C with primary anti-SARS-CoV-2 N antibody or anti-IAV NP antibody, followed by incubation with corresponding fluorophore-conjugated secondary antibody for 1 hour at room temperature. Nuclei were stained with 1× DAPI in PBS for 5 minutes. Finally, the coverslips were mounted onto glass slides using Prolong^TM^ Diamond Antifade Mountant. Fluorescence images were acquired using an epifluorescence microscope (Olympus IX71) equipped with a digital camera (Hamamatsu C13440) and a 20× objective lens. For each biological replicate, two to three random fields of view were imaged to assess the average infection rate.

#### Image Analysis and Infection Quantification

Fluorescence images were analyzed using CellProfiler software (v4.2.8). Primary objects (nuclei) were identified based on DAPI staining. Cell segmentation was performed using the Distance-N algorithm, where the cytoplasmic region was defined by expanding 10 pixels (approximately 3.18 μm) outward from each identified nucleus. The average fluorescent intensity of SARS-CoV-2 N or IAV NP signals was quantified within the segmented cytoplasmic area. The intensity threshold for determining whether a cell was infected was established based on the overall viral protein signal in the negative (mock) and positive (PBS) control groups for the infection. Cells with viral protein signal intensities exceeding this threshold were classified as infected, and the infection rate was calculated as the percentage of infected cells among the total number of cells, as determined by the number of segmented nuclei.

## Statistical Analysis

Statistical significance was determined by one-way analysis of variance (ANOVA) followed by Tukey’s post-hoc test. *p*-values < 0.05 were considered statistically significant (*p* > 0.05 [ns, not significant], *p* ≤ 0.05 [^∗^], *p* ≤ 0.01 [^∗∗^], *p* ≤ 0.001 [^∗∗∗^], *p* ≤ 0.0001 [^∗∗∗∗^]). All statistical analyses were performed using GraphPad Prism software (version 11) and OriginPro (2024).

## Acknowledgments

The authors acknowledge Dr. Syuan-Wei Li from Catalysis Facility for assistance with DLS and zeta potential measurements and Dr. Chun-Hong Yeh from NMR Core Facility for NMR measurements (Institute of Chemistry, Academia Sinica). The authors also thank Mr. Jing-Ting Zhu for rheological measurements, Mr. Arbaz Akhtar for virus-related experiments, and Dr. Gerhard Blankenburg for particle-tracking video acquisition during the development of Python-based analysis workflow (Institute of Physics, Academia Sinica); and Dr. Shih-Yin Ho and Prof. Wen-Chun Liu, from Infectious Disease Core Facility, Biomedical Translational Research Center (BioTrec), Academia Sinica, for performing the material biocompatibility assay. SARS-CoV-2 infection experiments were conducted in the Biosafety Level 3 Facility, Institute of Biomedical Sciences, Academia Sinica, with support from the Academia Sinica Core Facility and Innovative Instrument Project (Grant AS-CFII-115-A4). We acknowledge funding support from Academia Sinica (AS-IA-109-M04 and AS-iMATE-115-12) and National Science and Technology Council (NSTC 111-2112-M-001-029-MY3 and 114-2112-M-001-036-MY3). A.A. acknowledges fellowship support from the Nanoscience and Technology Program, Taiwan International Graduate Program (TIGP), Academia Sinica, Taiwan.

## Author contributions

A.A. conceived the study, designed the experimental work, performed the majority of the experiments, analyzed the data, and wrote the manuscript. H.Y.L and H.W.H. performed the virus-related experiments and analyzed the corresponding data, while A.A. contributed to the experimental design and preparation of the hydrogel formulations used in these studies. A.I.C. contributed to the synthesis, NMR characterization and rheological experiments, and participated in the interpretation and discussion of the results. J.H.W. contributed to the synthesis experiments and discussion of the experimental findings. I.H.W. contributed to the conceptualization and design of the virus-related experiments and to the interpretation and writing of the virology-related results. J.R.H. contributed to the interpretation and discussion of the rheological results. C.F.C. acquired the funding, supervised the project, provided overall scientific guidance, reviewed and revised the manuscript. All authors reviewed and approved the final version of the manuscript.

## Data Availability Statement

Data supporting the findings of this study are provided in the article and its Supporting Information. Additional data and python code are available from the corresponding author upon request.

## References

1. R. Bansil, B. S. Turner, "The biology of mucus: Composition, synthesis and organization," Advanced Drug Delivery Reviews 124(2018): 3–15.

2. A. McShane, J. Bath, A. M. Jaramillo, et al., "Mucus," Current Biology 31(2021): R938–R945.

3. M. Kesimer, C. Ehre, K. A. Burns, C. W. Davis, J. K. Sheehan, R. J. Pickles, "Molecular organization of the mucins and glycocalyx underlying mucus transport over mucosal surfaces of the airways," Mucosal Immunology 6(2013): 379–392.

4. M. A. McGuckin, S. K. Lindén, P. Sutton, T. H. Florin, "Mucin dynamics and enteric pathogens," Nature Reviews Microbiology 9(2011): 265–278.

5. D. J. Thornton, K. Rousseau, M. A. McGuckin, "Structure and function of the polymeric mucins in airways mucus," Annual Review of Physiology 70(2008): 459–486.

6. B. Button, H. P. Goodell, E. Atieh, et al., "Roles of mucus adhesion and cohesion in cough clearance," Proceedings of the National Academy of Sciences 115(2018): 12501–12506.

7. O. Lieleg, K. Ribbeck, "Biological hydrogels as selective diffusion barriers," Trends in Cell Biology 21(2011): 543–551.

8. B. Button, L.-H. Cai, C. Ehre, et al., "A periciliary brush promotes the lung health by separating the mucus layer from airway epithelia," Science 337(2012): 937–941.

9. A. G. Henderson, C. Ehre, B. Button, et al., "Cystic fibrosis airway secretions exhibit mucin hyperconcentration and increased osmotic pressure," The Journal of Clinical Investigation 124(2014): 3047–3060.

10. T. Kato, G. Radicioni, M. J. Papanikolas, et al., "Mucus concentration–dependent biophysical abnormalities unify submucosal gland and superficial airway dysfunction in cystic fibrosis," Science Advances 8(2022): eabm9718.

11. R. Bej, R. Haag, "Mucus-inspired dynamic hydrogels: synthesis and future perspectives," Journal of the American Chemical Society 144(2022): 20137–20152.

12. G. Ruiz-Pulido, D. I. Medina, "An overview of gastrointestinal mucus rheology under different pH conditions and introduction to pH-dependent rheological interactions with PLGA and chitosan nanoparticles," European Journal of Pharmaceutics and Biopharmaceutics 159(2021): 123–136.

13. M. Marczynski, K. Jiang, M. Blakeley, et al., "Structural alterations of mucins are associated with losses in functionality," Biomacromolecules 22(2021): 1600–1613.

14. J. P. Celli, B. S. Turner, N. H. Afdhal, et al., "Rheology of gastric mucin exhibits a pH-dependent sol− gel transition," Biomacromolecules 8(2007): 1580–1586.

15. L. Sardelli, D. P. Pacheco, A. Ziccarelli, et al., "Towards bioinspired in vitro models of intestinal mucus," RSC Advances 9(2019): 15887–15899.

16. C. E. Wagner, M. Krupkin, K. B. Smith-Dupont, et al., "Comparison of physicochemical properties of native mucus and reconstituted mucin gels," Biomacromolecules 24(2023): 628–639.

17. L. Esteban Enjuto, V. Taty Poaty, M. Bouveret, H. Song, S. Constant, J. Patarin, "Rheological comparison of sputum and reconstituted airway epithelium mucus," Scientific Reports 14(2024): 31660.

18. R. L. Howard, M. Markovetz, Y. Wang, et al., "Biochemical and rheological analysis of human colonic culture mucus reveals similarity to gut mucus," Biophysical Journal 120(2021): 5384–5394.

19. Y. Wang, R. Kim, C. Sims, N. Allbritton, "Building a thick mucus hydrogel layer to improve the physiological relevance of in vitro primary colonic epithelial models," Cellular and Molecular Gastroenterology and Hepatology 8(2019): 653–655. e655.

20. R. Bej, C. A. Stevens, C. Nie, et al., "Mucus-Inspired Self-Healing Hydrogels: A Protective Barrier for Cells against Viral Infection," Advanced Materials 36(2024): 2401745.

21. A. Mahalingam, J. I. Jay, K. Langheinrich, et al., "Inhibition of the transport of HIV in vitro using a pH-responsive synthetic mucin-like polymer system," Biomaterials 32(2011): 8343–8355.

22. S. Chandna, T. L. Povolotsky, C. Nie, et al., "Lignin-based mucus-mimicking antiviral hydrogels with enzyme stability and tunable porosity," ACS Applied Materials & Interfaces 17(2025): 8962–8975.

23. M. Kretschmer, R. Ceña-Diez, C. Butnarasu, et al., "Synthetic Mucin Gels with Self-Healing Properties Augment Lubricity and Inhibit HIV-1 and HSV-2 Transmission," Advanced Science 9(2022): 2203898.

24. M. Collado-González, M. Wu, A. Buckley, S. Berberolli, F. M. Goycoolea 2025, "Genipin crosslinks purified commercial pig gastric mucin to obtain a biomimetic mucus gel." in ChemRxiv.

25. K. Joyner, D. Song, R. F. Hawkins, R. D. Silcott, G. A. Duncan, "A rational approach to form disulfide linked mucin hydrogels," Soft Matter 15(2019): 9632–9639.

26. Z. Zhao, L. Chen, X. Li, et al., "Mucin-based injectable hydrogel loaded with Resolvin E1 for rectal mucosal repair after endoscopic submucosal dissection," Acta Biomaterialia 216(2026): 86-101.

27. R. Chen, K. Das, J. Feng, B. Thongrom, R. Haag, "Photo-regulated disulfide crosslinking: a versatile approach to construct mucus-inspired hydrogels," Chemical Science 16(2025): 5528–5537.

28. Q. Zhang, C.-Y. Shi, D.-H. Qu, Y.-T. Long, B. L. Feringa, H. Tian, "Exploring a naturally tailored small molecule for stretchable, self-healing, and adhesive supramolecular polymers," Science Advances 4(2018): eaat8192.

29. S. Asim, C. Tuftee, A. T. Qureshi, et al., "Multi-functional gelatin-dithiolane hydrogels for tissue engineering," Advanced Functional Materials 35(2025): 2407522.

30. B. R. Nelson, B. E. Kirkpatrick, C. E. Miksch, et al., "Photoinduced dithiolane crosslinking for multiresponsive dynamic hydrogels," Advanced Materials 36(2024): 2211209.

31. E. G. Plender, T. Prodanov, P. Hsieh, et al., "Structural and genetic diversity in the secreted mucins MUC5AC and MUC5B," The American Journal of Human Genetics 111(2024): 1700–1716.

32. F. L. Kearns, M. A. Rosenfeld, R. E. Amaro, "Breaking down the bottlebrush: atomically detailed structural dynamics of mucins," Journal of Chemical Information and Modeling 64(2024): 7949–7965.

33. A. Giri, M. Aquib, A. Choudhury, et al., "Lipoic Acid Based Redox-Responsive Degradable Antimicrobial Polymers," Macromolecular Rapid Communications 46(2025): e00224.

34. C. Wagner, K. Wheeler, K. Ribbeck, "Mucins and their role in shaping the functions of mucus barriers," Annual Review of Cell and Developmental biology 34(2018): 189–215.

35. S. Yuan, M. Hollinger, M. E. Lachowicz-Scroggins, et al., "Oxidation increases mucin polymer cross-links to stiffen airway mucus gels," Science Translational Medicine 7(2015): 276ra227–276ra227.

36. C. Kramer, H. Rulff, J. F. Ziegler, et al., "Ileal mucus viscoelastic properties differ in Crohn’s disease," Mucosal Immunology 17(2024): 713–722.

37. M. B. Stie, C. Cunha, Z. Huang, et al., "A head-to-head comparison of polymer interaction with mucin from porcine stomach and bovine submaxillary glands," Scientific Reports 14(2024): 21350.

38. T. Roe, T. Talbot, I. Terrington, et al., "Physiology and pathophysiology of mucus and mucolytic use in critically ill patients," Critical Care 29(2025): 68.

39. L. E. Morgan, A. M. Jaramillo, S. K. Shenoy, et al., "Disulfide disruption reverses mucus dysfunction in allergic airway disease," Nature Communications 12(2021): 249.

40. C. Taylor, A. Allen, P. W. Dettmar, J. P. Pearson, "The gel matrix of gastric mucus is maintained by a complex interplay of transient and nontransient associations," Biomacromolecules 4(2003): 922–927.

41. B. S. Schuster, J. S. Suk, G. F. Woodworth, J. Hanes, "Nanoparticle diffusion in respiratory mucus from humans without lung disease," Biomaterials 34(2013): 3439–3446.

42. O. Lieleg, I. Vladescu, K. Ribbeck, "Characterization of particle translocation through mucin hydrogels," Biophysical Journal 98(2010): 1782–1789.

43. H. M. Yildiz, C. A. McKelvey, P. J. Marsac, R. L. Carrier, "Size selectivity of intestinal mucus to diffusing particulates is dependent on surface chemistry and exposure to lipids," Journal of Drug Targeting 23(2015): 768–774.

44. B. C. Huck, O. Hartwig, A. Biehl, et al., "Macro-and microrheological properties of mucus surrogates in comparison to native intestinal and pulmonary mucus," Biomacromolecules 20(2019): 3504–3512.

45. R. Du, Q. Cui, L. Rong, "Competitive cooperation of hemagglutinin and neuraminidase during influenza A virus entry," Viruses 11(2019): 458.

46. S. Leach, "Release and breakdown of sialic acid from human salivary mucin and its role in the formation of dental plaque," Nature 199(1963): 486–487.

47. O. Lieleg, C. Lieleg, J. Bloom, C. B. Buck, K. Ribbeck, "Mucin biopolymers as broad-spectrum antiviral agents," Biomacromolecules 13(2012): 1724.

48. J. C. Crocker, D. G. Grier, "Methods of digital video microscopy for colloidal studies," Journal of Colloid and Interface Science 179(1996): 298–310.

49. M. K. Cheezum, W. F. Walker, W. H. Guilford, "Quantitative comparison of algorithms for tracking single fluorescent particles," Biophysical Journal 81(2001): 2378–2388.

50. K. I. Mortensen, L. S. Churchman, J. A. Spudich, H. Flyvbjerg, "Optimized localization analysis for single-molecule tracking and super-resolution microscopy," Nature Methods 7(2010): 377–381.

51. H. Qian, M. P. Sheetz, E. L. Elson, "Single particle tracking. Analysis of diffusion and flow in two-dimensional systems," Biophysical Journal 60(1991): 910–921.

52. X. Michalet, "Mean square displacement analysis of single-particle trajectories with localization error: Brownian motion in an isotropic medium," *Physical Review E—Statistical*, Nonlinear, and Soft Matter Physics 82(2010): 041914.

53. T. Savin, P. S. Doyle, "Static and dynamic errors in particle tracking microrheology," Biophysical Journal 88(2005): 623–638.

