## Supplementary Information for "Single-Component Photo-Crosslinked Lipoic Acid Functionalized Mucin Hydrogels: Dynamic Mucus Mimetics for Antiviral Protection"

### Table of Contents

1 Supplementary Results

1.1 Quantification of Accessible Thiol and Amine Groups in Commercial and Re-purified PGM (p. 3)

1.2 Large Amplitude Oscillatory Strain Sweeps of MUC-LA Hydrogel (p. 3)

1.3 CCK-8 Cytocompatibility Assay (p. 4)

1.4 Concentration Dependent Barrier Efficacy of Mucin Hydrogels Against Viral Transmission (p. 4)

2 Experimental Procedures

2.1 Materials and Methods (p. 5)

2.2 Re-purification of Commercial Porcine Gastric Mucin (p. 6)

2.3 Quantification of Accessible Amine and Thiol Groups in Commercial and Re-purified PGM (p. 6)

2.4 Particle Detection and Trajectory Analysis (p. 7)

2.5 Cytocompatibility of MUC-LA Polymer Solution and Hydrogels (p. 7)

2.6 Image Analysis and Infection Quantification with the Full-Slide Scanning System (p. 8)

3 Supplementary Figures

Figure S1-S6 (pp. 9-11)

4 Supplementary Videos

Video S1 and S2

References (p. 13)

### 1 Supplementary Results:

#### 1.1 Quantification of Accessible Thiol and Amine Groups in Commercial and Re-purified PGM

The availability of accessible thiol (SH) and primary amine ($NH_2$) groups in commercial and re-purified porcine gastric mucin (PGM) type II and III w evaluated to identify suitable functional groups for subsequent chemical modification **(Figure S1)**. Commercial PGM type II showed a higher level of accessible thiols than type III, whereas re-purification caused a marked decrease in thiol content for both mucin types **(Figure S1a)**. Because the DTNB assay detects accessible reduced sulfhydryl groups, the measured values represent thiols available for chemical reaction rather than the total cysteine content of mucin. The reduced thiol availability after re-purification therefore suggests that native sulfhydryl groups may be less suitable for controlled disulfide network formation. Structural and functional alterations of commercial mucins associated with processing have been reported previously.[1] Alternative thiol-sensitive reagents such as 4,4′-dithiodipyridine (DTDP) have also been used to quantify accessible thiols in mucin preparations.[1-2] In contrast, TNBSA analysis showed that a substantial number of primary amine groups remained accessible after re-purification **(Figure S1b)**. Although the accessible amine content decreased after re-purification, both PGM types II and III retained an estimated 100-150 amine groups per mucin molecule based on the representative molecular mass used for normalization. This difference between thiol and amine availability provided the basis for using primary amines as conjugation sites for lipoic acid (LA), thereby introducing a controlled dithiolane functionality into the mucin structure. PGM type II was selected for subsequent LA functionalization because it retained sufficient accessible primary amines after re-purification while also providing a more economical source for larger-scale hydrogel preparation

#### 1.2 Large Amplitude Oscillatory Strain Sweeps of MUC-LA Hydrogel

Large-amplitude oscillatory strain sweeps of the 5% (w/v) MUC-LA hydrogel were performed at a constant angular frequency of 10 rad $s^{-1}$ to determine the nonlinear deformation regime **(Figure S2).** At low strain, $G'$ remained substantially higher than $G''$, indicating a predominantly elastic response. With increasing strain, $G'$ gradually decreased while $G''$ increased, and the two moduli crossed at approximately 500% strain. This crossover indicates yielding of the hydrogel network and loss of the elastic-dominated response. Based on this result, 1000% strain was selected for the subsequent cyclic disruption and recovery measurements to ensure deformation well beyond the yielding regime

#### 1.3 CCK-8 Cytocompatibility Assay

The cytocompatibility of MUC-LA was evaluated by an indirect transwell-based exposure assay to determine whether soluble components released from the material affected MDCK cell viability. Leachates collected from photo-crosslinked hydrogels and non-crosslinked precursor solutions at 2%, 3%, and 5% (w/v) after 24, 48, and 72 h were assessed using the CCK-8 assay. For the crosslinked hydrogels, cell viability remained comparable to or above the mock control across all investigated concentrations and exposure periods **(Figure S5a)**, with no evident reduction in metabolic activity. This indicates that the photo-crosslinked MUC-LA formulations did not release appreciable levels of cytotoxic soluble components under the tested conditions. The non-crosslinked MUC-LA precursor solutions similarly showed no evident reduction in cell viability at 2%, 3%, or 5% **(Figure S5b),** although elevated CCK-8 responses were observed for some of the higher-concentration conditions. As CCK-8 measures cellular metabolic activity, responses above the normalized control were interpreted as maintained or elevated metabolic activity rather than direct evidence of increased cell proliferation. Previous work by Lieleg[3] showed that reconstituted porcine gastric mucin at 1% (w/v) did not compromise the viability of HeLa cells, supporting the intrinsic cytocompatibility of mucin as a biomaterial. Overall, these observations suggest that neither the precursor solutions nor the photo-crosslinked MUC-LA hydrogels produced an evident reduction in MDCK cell viability under the tested conditions.

#### 1.4 Concentration-Dependent Barrier Efficacy of Mucin Hydrogels Against Viral Transmission

To briefly explore the effect of mucin concentration on the barrier efficacy of the hydrogel, we evaluated a lower concentration 2% (w/v) and compared its ability to block SARS-CoV-2 transmission with that of 3% (w/v) mucin hydrogel in the transwell-based infection assay (**Figure S6)**. When 2% and 3% non-crosslinked mucin were applied directly, cell infection rates showed no significant difference. However, while the 3% mucin hydrogel completely impeded viral transmission, sporadic infected cells were observed in the transwell-based infection system employing the 2% mucin hydrogel. Using a full-slide scanning system for comprehensive cell analysis across the entire well, we determined the infection rate to be approximately 2.27% for the 2% mucin hydrogel group, accompanied by a distinct infection focus, whereas the 3% mucin hydrogel reduced the infection rate to 0%. We speculate that varying the mucin concentration alters the mesh network architecture of the hydrogel, leading to differential performances of these barriers.

### 2 Experimental Procedures:

#### 2.1 Materials and Methods

Porcine gastric mucin (PGM; Type II, bound sialic acid ≤ 1.2% and Type III, partially purified, bound sialic acid 0.5%-1.5%), bovine submaxillary mucin (BSM, M3895), Sodium phosphate dibasic, sodium phosphate monobasic, 5,5-dithio-bis-(2-nitrobenzoic acid, DTNB), 2,4,6-trinitrobenzene sulfonic acid (TNBSA), Sodium bicarbonate , L-lysine, α-Lipoic acid (LA), N-hydroxysuccinimide (NHS), N-(3-Dimethylaminopropyl)-N′-ethylcarbodiimide hydrochloride (EDC), 2-mercaptoethanol, sodium chloride, deuterium oxide (D2O) and fluorescein sodium salt were purchased from Sigma-Aldrich (St. Louis, MO, USA). L-Cysteine hydrochloride monohydrate, Glutathione (GSH), 2-(morpholin-4-yl) ethane-1-sulfonic acid hydrate (MES), FluoSpheres$^{TM}$ size kit#2 carboxylate-modified microsphere (F8888), EDTA and methylene blue were purchased from Thermo Scientific (USA). Hydrochloride (HCl), sodium dodecyl sulfate (SDS) and Dimethyl sulfoxide (DMSO) were purchased from J.T. Baker (USA). Spectra/Por® 4 dialysis membranes (12-14 kDa MWCO) were purchased from Repligen (USA). Methoxy-poly (ethylene glycol) mPEG-NH2, 5k was purchased from biopharma PEG (USA).

**For cell-based studies**, 24-well Transwell inserts with 0.4 µm pore size, (Corning Cat. No. 3470), and Cell Counting Kit-8 (CCK-8; TOOLS, Cat. No. TEN-CCK8) were used. Minimum Essential Medium (MEM) and Opti-MEM was purchased from Gibco (Thermo Fisher Scientific, Waltham, MA, USA). Fetal bovine serum (FBS) was purchased from Cytiva (Marlborough, MA, USA). Phosphate-buffered saline (PBS) was purchased from Protech Technology Enterprise Co. (Taipei, Taiwan). Bovine serum albumin (BSA) and goat serum were purchased from Bovogen Biologicals (East Keilor, VIC, Australia). Penicillin-streptomycin (P/S) was purchased from GeneTeks Biosciences (Taipei, Taiwan). Paraformaldehyde (PFA), glycine, Triton X-100, and DAPI were purchased from Sigma-Aldrich (St. Louis, MO, USA). Rabbit anti-SARS-CoV-2 nucleocapsid (N) antibody (GTX135357) and rabbit anti-IAV nucleoprotein (NP) antibody (GTX125989) were purchased from GeneTex, Inc. (Irvine, CA, USA). Alexa Fluor$^{TM}$ 568-conjugated goat anti-rabbit

IgG secondary antibody (A-11036) and Prolong™ Diamond Antifade Mountant were purchased from Invitrogen (Thermo Fisher Scientific, Waltham, MA, USA).

#### 2.2 Re-Purification of Commercial Porcine Gastric Mucin

The commercially available porcine gastric mucin (PGM) Types II and III were used and partially re-purified by following the previously reported protocol,[4] with slight modifications to remove low molecular weight impurities and residual salts. This step provides a cleaner and more reproducible starting material prior to functionalization. Briefly, the obtained mucin powder was dissolved in deionized (DI) water to prepare a 1% solution and stirred overnight at 37 °C to ensure complete hydration. The solution was then centrifuged at 2000 rpm for 20 minutes to remove large aggregates and insoluble impurities. The resulting supernatant was collected and subjected to dialysis using a 12-14 kDa (MWCO) membrane for three days at 4 °C with three exchanges of DI water daily to remove salts and low molecular weight contaminants. After dialysis, the re-purified mucin solution was freeze-dried and later stored at -20 °C until further use.

#### 2.3 Quantification of Accessible Amine and Thiol Groups in Commercial and Re-purified PGM

Accessible primary amines and free thiol groups in commercial and re-purified PGM (Types II and III) were quantified using 2,4,6-trinitrobenzenesulfonic acid (TNBSA) and Ellman's assays, respectively. Primary amine content was determined using the TNBSA assay according to a previously reported method[5] and the manufacturer's protocol, with minor modifications (Thermo Scientific, Manual D00386, TS-28997). L-Lysine (MW = 146.19 g $mol^{-1}$) was used to generate the calibration curve. Mucin samples were prepared at 100 μg $mL^{-1}$ and reacted with 0.01% (w/v) TNBSA in 0.1 M sodium bicarbonate buffer (pH 8.5). Aliquots (200 μL) were transferred to a 96-well plate and incubated at 37 °C for 2 h. The reaction was terminated by sequential addition of 10% (w/v) SDS and 1 N HCl, and the absorbance was measured at 335 nm using a microplate reader (Multiskan GO, Thermo Scientific, Finland). Measurements were performed in triplicate, and the accessible amine content was determined from the L-lysine calibration curve.

Free thiol groups were quantified using Ellman's reagent (5,5′-dithiobis(2-nitrobenzoic acid), DTNB) according to the established Ellman method,[6] and the manufacturer's protocol, with minor modifications (Thermo Scientific, MAN0011216_Ellmans_Reag_UG, 22582). Mucin samples were prepared at 1 mg $mL^{-1}$ in reaction buffer consisting of 0.1 M sodium phosphate (pH

8.0) containing 1 mM EDTA. L-Cysteine hydrochloride (Cys-HCl; MW = 175.6 g $mol^{-1}$) was used to construct the calibration curve. Samples (250 uL) were incubated with 50 μL of DTNB solution (4 mg $mL^{-1}$ in reaction buffer) for 30 min in a 96-well plate, followed by measurement of absorbance at 412 nm. Measurements were performed in triplicate, and thiol concentrations were determined from the Cys-HCl calibration curve. Accessible amine and thiol contents were calculated as μmol $mg^{-1}$ of mucin. To estimate the number of reactive groups per mucin molecule, the experimentally determined values were additionally normalized using a representative mucin molecular mass of ≈640 kDa while accounting for the intrinsic heterogeneity and polydispersity of mucin.

#### 2.4 Particle Detection and Trajectory Analysis

Particle candidates were preliminarily detected and linked using TrackPy with a feature diameter of 9 pixels, minimum separation of 7 pixels, search range of 6 pixels, memory of 2 frames and a preliminary minimum trajectory length of 20 frames. Minimum mass thresholds were 2500 and 7500 for the 100 and 200 nm particles, respectively. Particle trajectories were analyzed using the original video frame numbers so that displacement pairs were formed only between coordinates separated by the exact requested frame lag. MSD values were retained only when supported by at least 250 valid displacement pairs. Only positive localization-corrected MSD values were included for logarithmic power-law fitting.

#### 2.5 Cytocompatibility of MUC-LA Polymer Solution and Hydrogels

To evaluate the cytocompatibility of MUC-LA hydrogel, a 24-well transwell system with 0.4 μm pore size inserts (Corning, Cat. No. 3470) was used. Briefly, $1\times10^5$ MDCK cells/well were seeded into the lower chambers of a 24-well transwell plate. The following day, MUC-LA precursor solutions (2%, 3% and 5% w/v) were prepared in 1xPBS. A 50 μL aliquot of each precursor was loaded into transwell inserts under sterile conditions and UV-cured (365nm, 20 mW $cm^{-2}$) to form a hydrogel. The inserts were then transferred into the MDCK cell-containing wells, and an additional 50 μL of Opti-MEM was added to the apical side of both hydrogel and solution conditions. At 24, 48, and 72 h, 100 μL of conditioned medium was collected from the lower chamber and replaced with an equal volume of fresh Opti-MEM. The collected samples were stored at -80 °C. For cell viability measurements, MDCK cells were seeded in a 96-well plate at 2

$\times 10^4$ cells/well density at 24 h prior to the assay. The culture medium was aspirated and replaced with 100 μL of the collected hydrogel-treated medium from each time point. After 24 h incubation, cell viability was measured using the CCK-8 assay (CCK-8, TOOLS, Cat. No. TEN-CCK8) according to the manufacturer's instructions. Relative cell viability (%) was calculated by normalization of absorbance values recorded at 450 nm using Microplate reader (SpectraMax i3x, Molecular Devices) against untreated controls.

##### 2.6 Image Analysis and Infection Quantification with the Full-Slide Scanning System

Fluorescence images were acquired using an ImageXpress HCS.ai CONFOCAL High-Content Screening System (Molecular Devices, USA). To avoid edge-associated background artifacts, region-of-interest (ROI) imaging was selectively restricted to the central area of each cover-glass, strictly excluding the peripheral boundaries. Then, quantitative image analysis was performed using IN Carta Image Analysis Software (v2.9, Molecular Devices). Total cell number were determined by segmenting individual nuclei based on DAPI signals. Cell boundaries were subsequently delineated to quantify the mean fluorescence intensity of the SARS-CoV-2 N protein per cell. An intensity threshold for defining infection was established based on the negative (mock) and positive (PBS) control groups. Cells exhibiting N protein signal intensities above this threshold were classified as infected. The infection rate was calculated as the percentage of infected cells relative to the total number of cells within the analyzed ROIs.

### 3 Supplementary Figures:

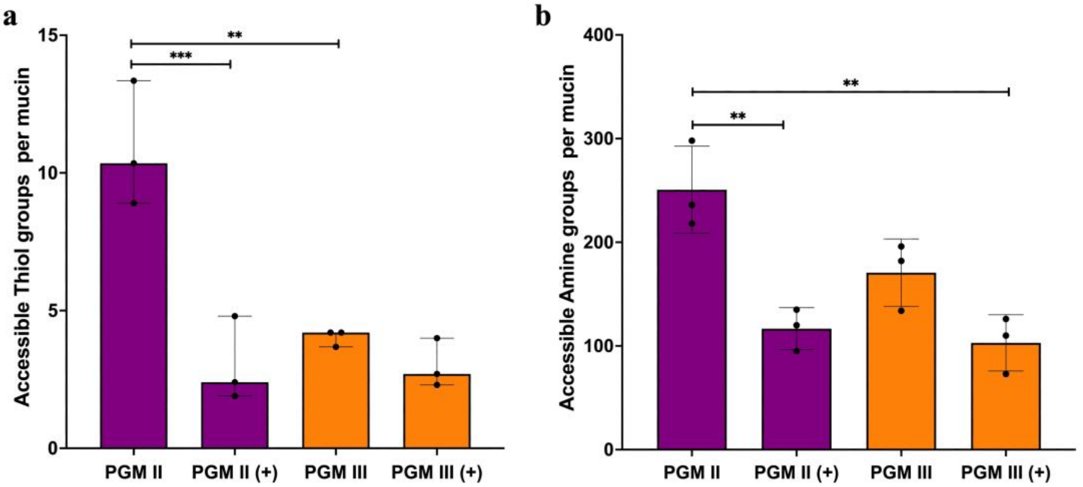


**Figure S1.** Quantification of accessible functional groups in commercial and re-purified PGM mucins. (a) Accessible thiol (-SH) groups and (b) accessible amine (-$NH_2$) groups per mucin molecule for PGM Types II and III before and after re-purification. (+) indicates re-purified mucin. Data are presented as mean ± [SD], n = [3].

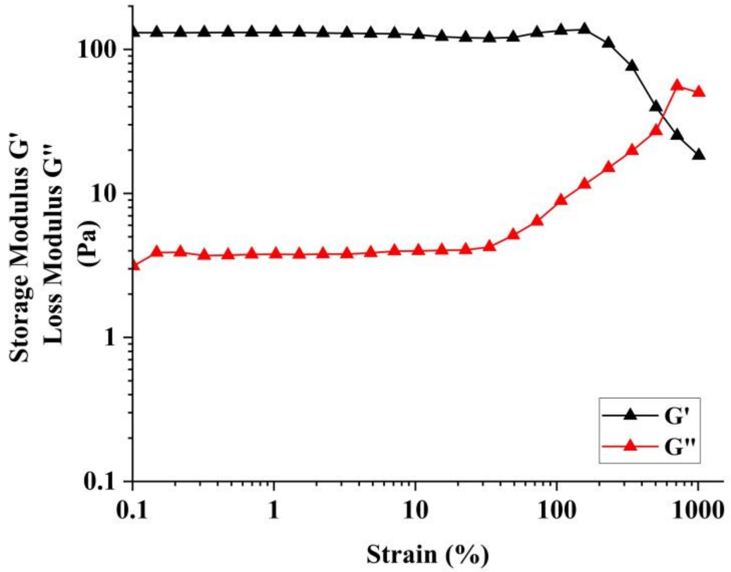


**Figure S2.** Large-amplitude oscillatory strain sweeps of the 5% MUC-LA hydrogel at 10 rad $s^{-1}$, showing the crossover of G′ and G″ at high strain and the loss of elastic dominant response.

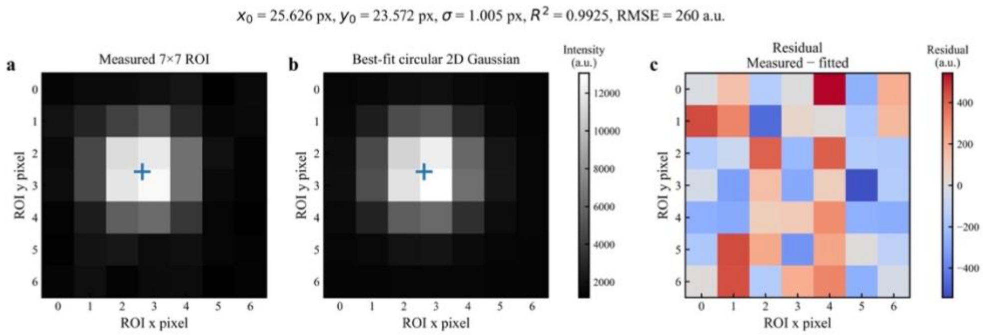


**Figure S3.** Validation of centrosymmetric two-dimensional Gaussian localization. a) Representative measured 7 × 7-pixel fluorescence region of interest (ROI) containing a 200 nm PEG-coated particle, with the fitted subpixel centroid indicated by the blue cross. b) Corresponding best-fit centrosymmetric two-dimensional Gaussian intensity distribution. c) Residual image calculated as $I_{\text{measured}} - I_{\text{fitted}}$ showing the pixel-wise difference between the measured and fitted fluorescence intensities. For the representative localization shown $x_0$ = 25.626px, $y_0$ = 23.572px, $\sigma$ = 1.005px, $R^2$ = 0.9925 and RMSE = 260 a.u. The small residual magnitude relative to the particle signal supports the suitability of the circular Gaussian model for subpixel centroid localization.

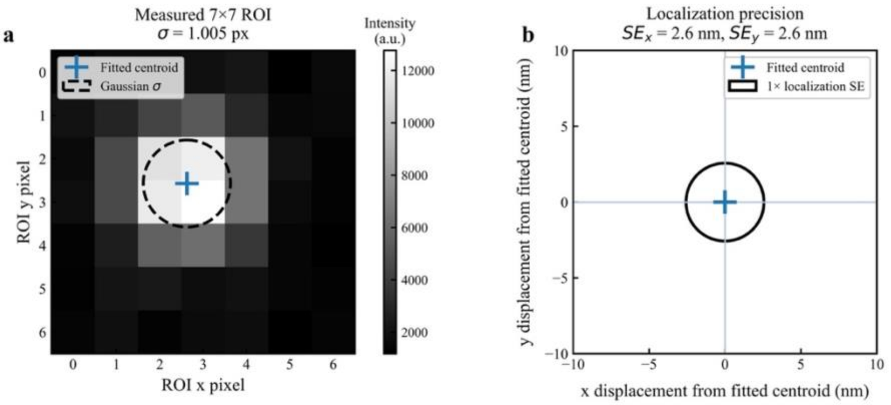


**Figure S4.** Gaussian spot width and subpixel localization precision. a) Representative measured 7 × 7-pixel fluorescence ROI containing a 200 nm PEG-coated particle, with the fitted centroid indicated by the blue cross. The dashed circle represents the fitted circular Gaussian width, $\sigma$ = 1.005px corresponding to approximately 160 nm. b) Localization precision plotted relative to the fitted centroid, defined as the origin. The black contour represents 1× localization standard error, with $SE_x$ = 2.6 nm and $SE_y$ = 2.6 nm. The Gaussian width describes the spatial extent of the fluorescence intensity distribution, whereas the localization standard error represents the uncertainty of the fitted centroid position.

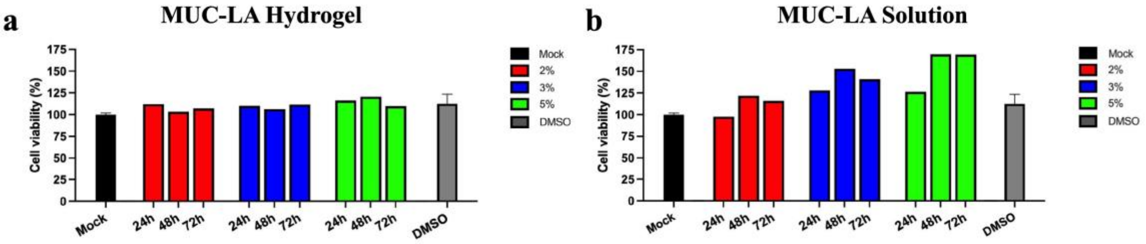


**Figure S5.** Cytocompatibility assessment of MUC-LA formulations using MDCK cells. (a) Relative cell viability following exposure to leachates collected from photo-crosslinked MUC-LA hydrogels (2%, 3%, and 5% w/v) after 24, 48, and 72 h. (b) Relative cell viability following exposure to leachates from the corresponding non-crosslinked MUC-LA precursor solutions. Cells viability was determined by CCK-8 assay and normalized to the mock control. Mock and DMSO controls are presented as mean ± SD (n = 3), whereas MUC-LA treatment conditions represent single measurements (n = 1) and are shown without error bars.

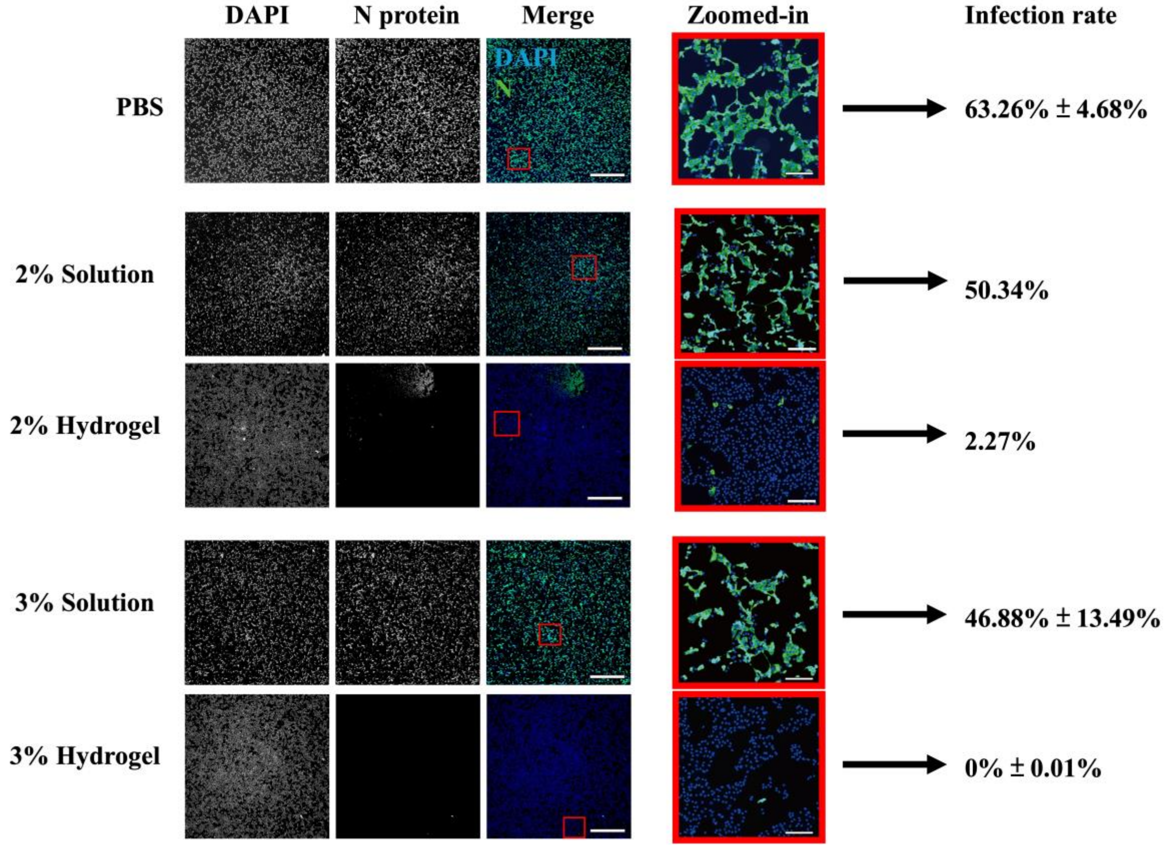


**Figure S6.** Concentration-dependent barrier efficacy of MUC-LA solution and hydrogels against viral transmission. Immunofluorescence analysis of SARS-CoV-2 infection in Vero E6 cells in the transwell-based infection system at 24 hours post-inoculation. The transwell was layered with PBS, MUC-LA solution (2% or 3%), or hydrogel (2% or 3%). Viral nucleocapsid (N) protein is shown in green, and cell nuclei counterstained with DAPI are shown in blue. The images were acquired using a full-slide scanner. Scale bar = 2000 μm (Zoomed-in images, 200 μm).

#### 4 Supplementary Videos

**Video S1:** Macroscopic Flow Behavior of 3% MUC-LA Hydrogel

The video shows the macroscopic deformation and flow behavior of the 3% MUC-LA hydrogel during manual handling, illustrating its soft viscoelastic behavior.

**Video S2:** Size-Dependent Motion of 100 and 200 nm PEGylated Particles in 3% MUC-LA Hydrogel

The video shows representative side-by-side recordings of 100 and 200 nm PEGylated particles within the 3% hydrogel. The 100 nm particles exhibit greater spatial displacement, whereas the 200 nm particles show more confined motion, consistent with the size-dependent transport restriction behavior quantified by particle-tracking analysis. The recordings are displayed at accelerated playback speed (25 fps), with a 5 µm scale bar shown for spatial reference.

**References:**


1. M. Marczynski, K. Jiang, M. Blakeley, et al., "Structural alterations of mucins are associated with losses in functionality," *Biomacromolecules* 22(2021): 1600-1613.
2. C. K. Riener, G. Kada, H. J. Gruber, "Quick measurement of protein sulfhydryls with Ellman's reagent and with 4, 4′-dithiodipyridine," *Analytical and Bioanalytical Chemistry* 373(2002): 266-276.
3. O. Lieleg, C. Lieleg, J. Bloom, C. B. Buck, K. Ribbeck, "Mucin biopolymers as broad-spectrum antiviral agents," *Biomacromolecules* 13(2012): 1724.
4. C. E. Wagner, M. Krupkin, K. B. Smith-Dupont, et al., "Comparison of physicochemical properties of native mucus and reconstituted mucin gels," *Biomacromolecules* 24(2023): 628-639.
5. P. Cayot, G. Tainturier, "The quantification of protein amino groups by the trinitrobenzenesulfonic acid method: a reexamination," *Analytical Biochemistry* 249(1997): 184-200.
6. G. L. Ellman, "Tissue sulfhydryl groups," *Archives of Biochemistry and Biophysics* 82(1959): 70-77.
